# Characterization of core bacterial species in the *Daphnia magna* microbiota using shotgun metagenomics

**DOI:** 10.1101/645630

**Authors:** Reilly O. Cooper, Clayton E. Cressler

## Abstract

**Background:** The keystone zooplankton *Daphnia magna* has recently been used as a model system for understanding host-microbiota interactions. However, the bacterial species present and functions associated with their genomes are not well understood. In order to understand potential functions of these species, we combined 16S rRNA sequencing and shotgun metagenomics to characterize the whole-organism microbiota of *Daphnia magna*.

**Results:** Five metagenome-assembled genomes (MAGs) were assembled from the *Daphnia magna* microbiota. Phylogenetic placement of these MAGs indicated that two belong to the *Limnohabitans* genus, one to *Polaromonas*, one to *Pedobacter*, and one unclassifiable below the Burkholderiaceae family. Average nucleotide identity of these MAGs to their closest sequenced relative was <95%, suggesting these may be new species in known genera. 16S rRNA community profiling shows that the *Daphnia magna* microbiota is distinct from its culture environment. Genes involved in host colonization and immune system evasion were detected across the MAGs. Some metabolic pathways were specific to some MAGs, including sulfur oxidation, nitrate reduction, and flagellar assembly. Threonine and arginine exporters were encoded by the *Limnohabitans* and Burkholderiaceae MAGs, and pathways for key vitamin biosynthesis and export were identified across MAGs.

**Conclusions:** In this study, we characterize five metagenome-assembled bacterial genomes within the *Daphnia magna* microbiota. Our examination of functions associated with these genomes shows a diversity of nutrient acquisition and metabolism pathways present that may benefit the host, as well as genomic signatures of host association and immune system evasion.

## Introduction

Organisms are hosts to complex communities of microorganisms that live on any tissue in contact with the environment, collectively known as the microbiota. Species in the microbiota may provide beneficial functions to the host and to other species in the microbiota, including nutrient acquisition and uptake [1], production of host-accessible metabolites [2], host immune system priming [3], and direct pathogen protection [4]. However, characterizing these beneficial host-microbe and microbe-microbe interactions in biologically relevant systems can be difficult due to the number of core species present in the microbiota and the variation in an individual organism’s microbiota over time from dietary changes or differential environmental exposure [5].

The zooplankton *Daphnia magna* provides a useful model for studying functional relationships between microbes and their hosts. *Daphnia* species are used as a model system in ecology, ecotoxicology, and host-parasite dynamics due to their well-documented life cycle and rapid asexual reproduction [6]. The ability to raise *Daphnia magna* clonally in the laboratory allows for genetically identical hosts to be used experimentally, reducing the impact of genetic variation on the microbiota. Furthermore, their indiscriminate filter feeding within the water column allows for control over food input. *Daphnia* are colonized with bacteria throughout their entire body cavity and gut [7,8]. Composition of the *Daphnia* microbiota appears to be similar in spatially unique populations [8,9], suggesting mechanisms of acquisition and cultivation of these microbes by the host. The *Daphnia* microbiota is relatively simple at the class level, with β-proteobacteria, γ-proteobacteria, and Flavobacteriia consistently identified at high relative abundances [9,10]. While some work on the *Daphnia* microbiota has used shotgun sequences to examine potential bacterial symbionts [7], the majority of studies have used coarse-level 16S rRNA analyses to profile the taxa present. Species-level identification and subsequent functional profiling of the *Daphnia* microbiota has not been achieved with these studies, as resolution of the microbiota using 16S rRNA sequencing can only reliably identify bacterial genera. Much of the genomic content of *Daphnia*-associated bacteria is unknown, and potentially novel taxa may not be identified by standard 16S rRNA techniques.

Beyond identifying the composition of the microbiota, it is clear that this composition affects host fitness. The microbiota as a whole has been implicated in nutrient acquisition and breakdown of toxic compounds [11], and survival and growth are affected by the presence of, and perturbations to, the microbiota [12–14]. Host fecundity has been specifically tied to the most abundant genus in the *Daphnia* microbiota, *Limnohabitans*. Entirely bacteria-free *Daphnia* experience significant declines in fecundity, but monocolonization of bacteria-free *Daphnia* with *Limnohabitans* restores fecundity to that of *Daphnia* with a complete microbiota [15]. However, it is unclear what metabolic functions *Limnohabitans* provides to achieve these effects. How the other species in the *Daphnia* microbiota contribute to host life history and what functions these species may be providing is entirely unclear.

Here, we use shotgun metagenomics to characterize the bacterial species present in the *Daphnia magna* microbiota. The metagenome-assembled genomes generated from this data were then used to examine potential metabolic functions of the microbiota in total and for each species assembled. This study is the first to report metagenome-assembled genomes of bacteria in the *Daphnia magna* microbiota and is the first to suggest functions based on gene content of these microbes. This increased resolution allows us to formulate testable hypotheses about the metabolic interactions happening between the host and microbes, and among microbes, that impact host fitness. This functional knowledge will provide a new lens for studying this important ecological model system.

## Results

### Shotgun sequencing, assembly, and binning

Shotgun sequencing of the adult and juvenile *Daphnia magna* microbiota resulted in 20.19Mbp of paired-end Illumina read data, which was then reduced to 9.64Mbp after quality trimming and host genome filtering. A co-assembly of all four samples generated 174,991 contigs (N50 of 1,923bp; longest assembled contig 226,522bp). Summaries of assembly statistics for the co-assemblies and the individual sample assemblies can be found in Table 1. Identification of high-quality reads and ≥1000bp assembled contigs using Kraken, Kaiju, and MetaPhlAn2 indicated 18 genera that were present as more than 1% of the sample (Supplementary Figure 1). Of the 18 genera, only *Limnohabitans* was identified by all three tools in reads and contigs. Other abundant genera included *Pedobacter*, *Flavobacterium*, *Polaromonas* and other unclassified Burkholderiaceae. 16S rRNA sequencing of the library preparation and DNA extraction kits resulted in <50 reads (Figure 1A). 16S rRNA community profiles of the *Daphnia magna* food source, *Chlamydomonas reinhardtii*, the COMBO culture media *Daphnia magna* are raised in, and samples of 5 healthy adult *Daphnia magna* were found to have differences in composition (Figure 1A). The *Chlamydomonas* samples showed reduced relative abundance of Proteobacteria as compared to adult *Daphnia* and the culture media. The culture media showed higher relative abundance of Actinobacteria, and healthy *Daphnia magna* primarily were colonized by Proteobacteria and Bacteroidetes. These same community profiles were analyzed using an unweighted principal coordinate analysis (PCoA) on the unweighted UniFrac distances between samples, which represents the phylogenetic relatedness between samples based on ASV presence and absence (Supplementary Figure 2). The three sample groups were found to cluster separately, with no overlap.

**Figure 1.**
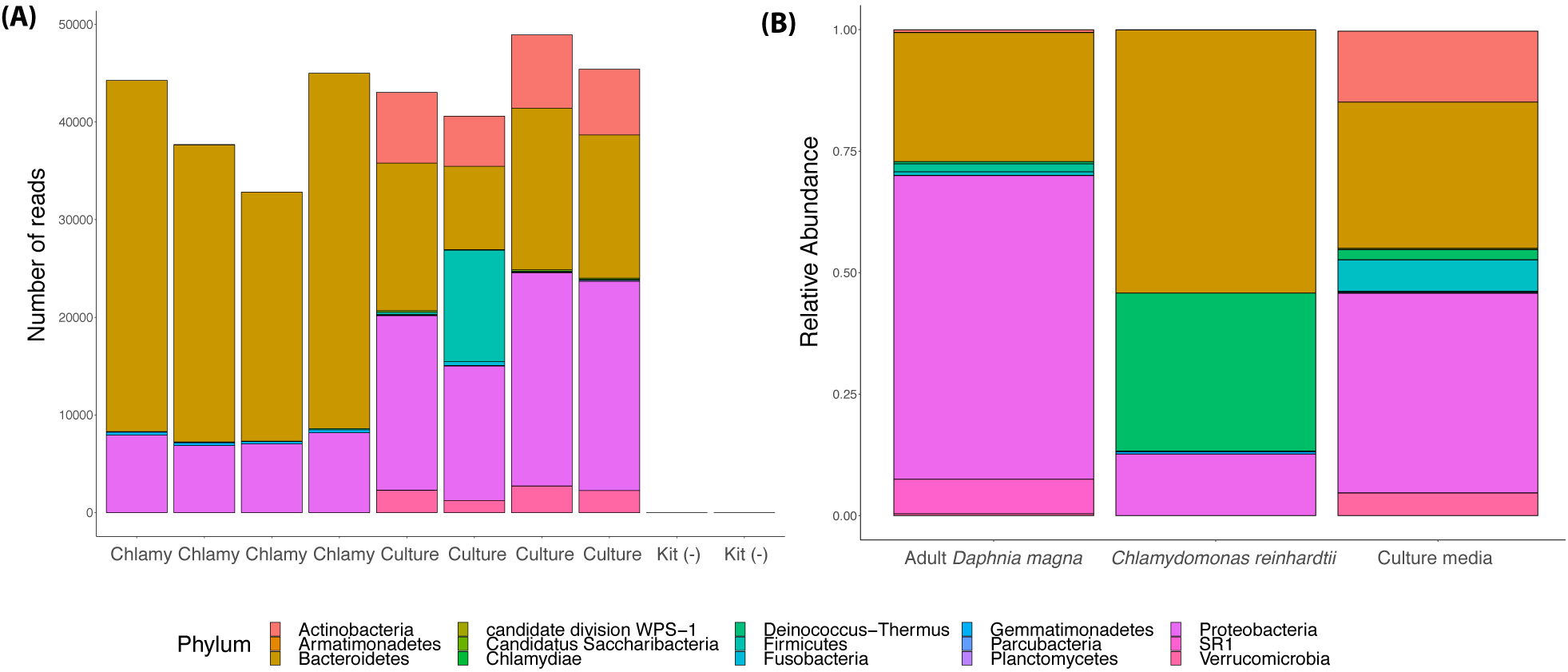
**(A)** Phylum-level 16S rRNA profiles of *Chlamydomonas reinhardtii* (Chlamy, n = 4), COMBO culture media (Culture, n= 4), and the DNA extraction kit and library preparation kit used for sequencing (Kit (-), n=2). **(B)** Relative abundance of phyla generated from 16S rRNA community profiles in *Chlamydomonas reinhardtii*, healthy adult *Daphnia magna*, and the culture media (all n=4).

**Table 1.**
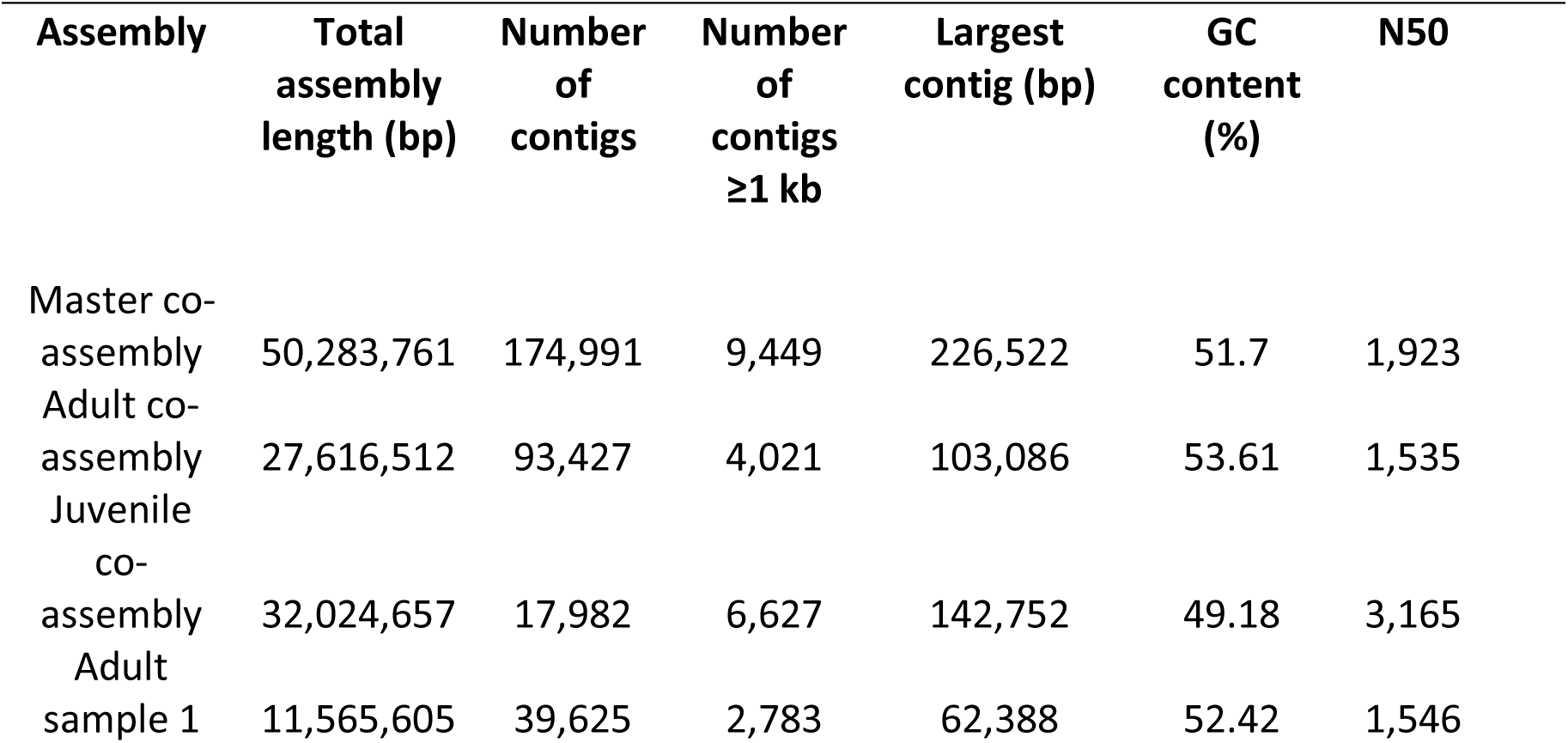
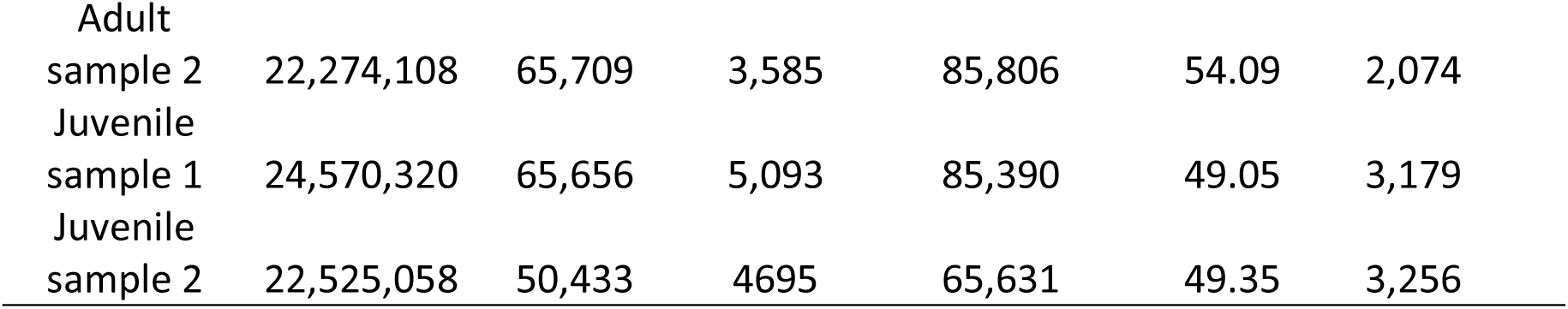
Summary of assembly statistics for individual sample assemblies and co-assemblies of all four samples (master co-assembly), the two adult samples (adult co assembly), and the two juvenile samples (juvenile co-assembly).

CONCOCT clustering of contigs from the master co-assembly by tetranucleotide composition, GC content, and coverage resulted in 19 bins of draft and incomplete genomes. Manual curation, refinement, and merging using Anvi’o resulted in 15 bins, with 7 meeting the ≥50 quality metric. The adult-only co-assembly generated 12 bins, which further refined to 5 bins, with 4 meeting the ≥50 quality metric (Supplementary Figure 5). Three of the four high-quality bins were also present in the master co-assembly and one unique bin was identified. The juvenile-only co-assembly generated 15 bins, which further refined to 7 bins, with 4 meeting the ≥50 quality metric (Supplementary Figure 6). Because the master co-assembly generated bins with higher coverage or higher completeness, we used GTDB-Tk on bins refined in the master co-assembly only. Phylogenetic placement using GTDB-Tk for the seven high- and medium-quality bacterial MAGs identified from the master co-assembly placed two in the *Limnohabitans* genus (*Limnohabitans* MAG 1 and MAG 2), one in the *Polaromonas* genus (*Polaromonas* MAG), one unable to be placed below the Burkholderiaceae family (Burkholderiaceae MAG), one in the *Pedobacter* genus (*Pedobacter* MAG), one in the *Emticicia* genus, and one unable to be placed below the Chitinophagales order (Supplementary Table 1; Supplementary Figure 4). None of the MAGs mapped to previously identified species in the GTDB-Tk database. Average nucleotide identity of the two *Limnohabitans* MAGs was measured at 86.6% similarity. This is less than the 95% cutoff generally accepted for strain-levels similarity, suggesting the MAGs are likely separate species [16].

To examine potential functions of the Daphnia magna microbiota, we focused on the three most complete MAGs: unknown Burkholderiaceae, 99.28% complete, 11.5X mean coverage; *Pedobacter* sp., 98.56% complete, 14.87X mean coverage; and *Polaromonas* sp., 82.73% complete, 15.3X mean coverage. We also examined the two *Limnohabitans* MAGs: sp1, 78.42% complete, 29.99X coverage; sp2, 60.43% complete, 14.16X coverage (Figure 2). Though the *Limnohabitans* MAGs were not over 90% complete, according to read-based and contig-based identification tools they were the most abundant, and coverage of both exceeded 30x in the juvenile *D. magna* samples. Furthermore, prior work has indicated the presence and importance of this genus in the *Daphnia magna* microbiota, suggesting unique functions may be present in these genomes. We extracted the nearest matching reference genome for each of the five MAGs above and calculated ANI for each to further investigate if these MAGs were novel or just new strains of already sequenced species (Table 2).

**Figure 2.**
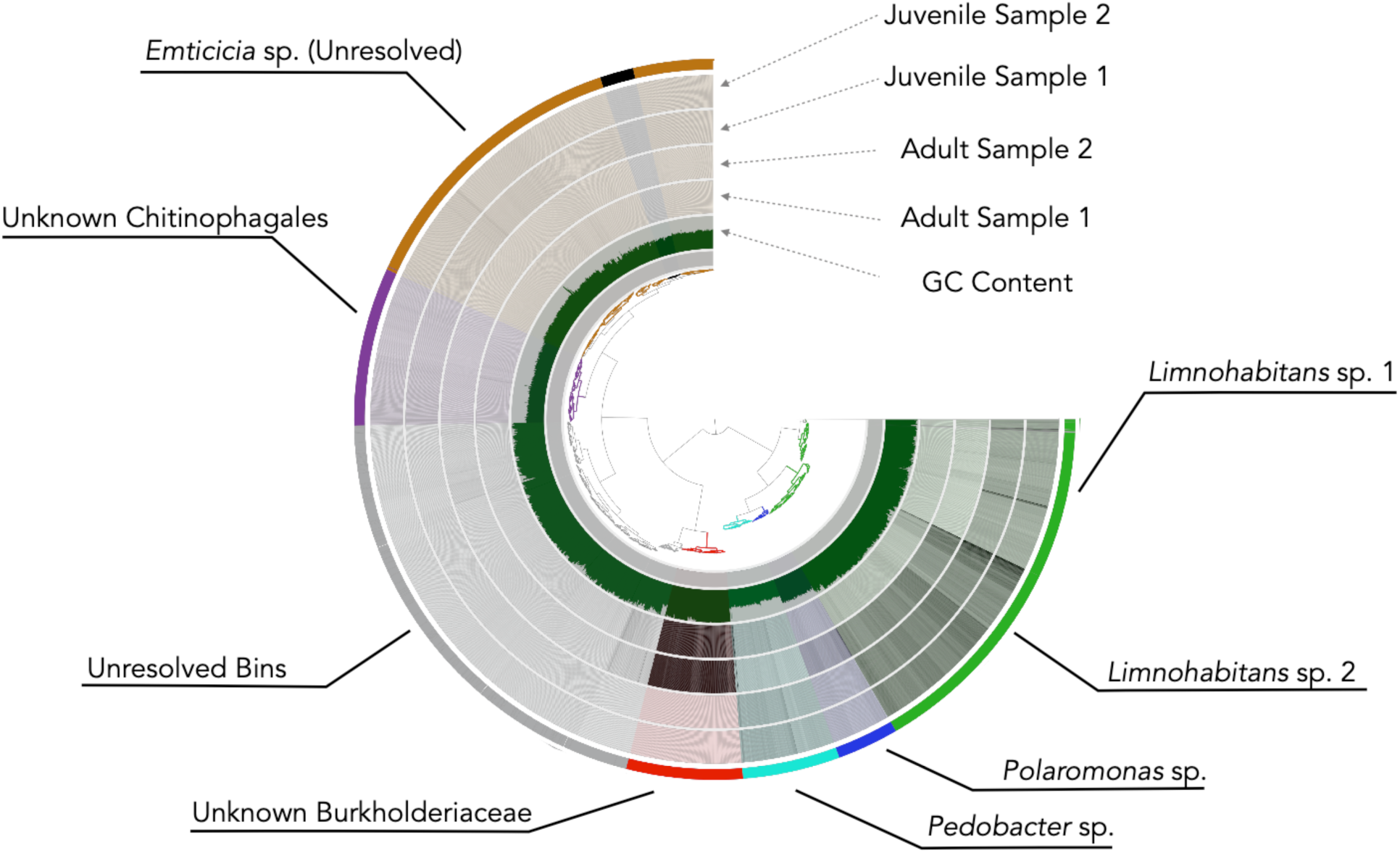
**(A)** Anvi’o display of the two adult and two juvenile samples with the metagenome-assembled genomes highlighted in color. The innermost dendogram represents similarity among contigs based on sequence composition, GC content (green inner ring), and differential coverage (black bars across the four grey rings). The outermost layer shows the genome bins. Genome bin identification performed by GTDB-Tk.

**Table 2.**
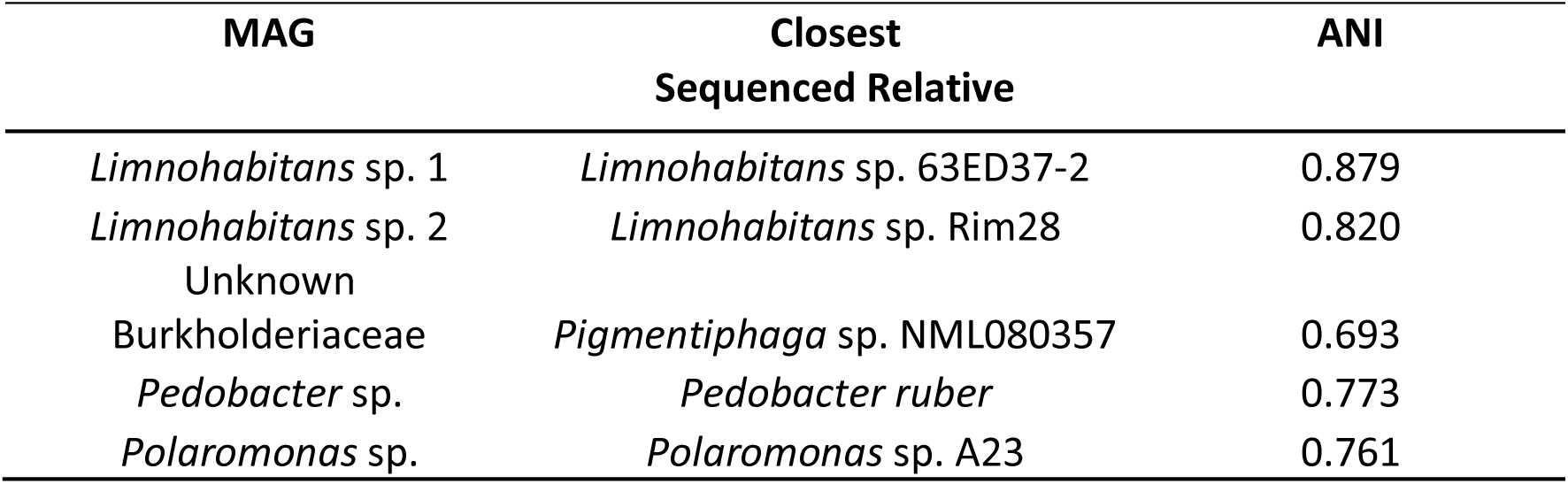
Average nucleotide identity of the five MAGs of interest to their nearest sequenced relative. The closest sequenced relative was identified from placement within GTDB-Tk’s reference phylogeny output.

### Functional profiling of the five high- and medium-quality bacterial metagenome-assembled genomes

Contigs from the entire metagenome and from the 5 MAGs independently were analyzed for potential coding sequences (CDS) using Prokka. All identified CDS were then queried against the KEGG database using GhostKOALA and identified orthologs of functional genes were grouped and quantified into KEGG functional categories (Figure 3). We focused on pathways associated with respiration, carbohydrate metabolism, amino acid metabolism, other energy metabolism, and transport. These functional categories indicate not only general metabolism for these species, but also could indicate what functions these species are providing to the host. Out of the 7,453 CDS from the five high-quality MAGs that mapped to KEGG pathways, 707 (9.48%) were associated with carbohydrate metabolism, 642 (8.71%) were associated with amino acid metabolism, and 358 (4.80%) were associated with other energy metabolism (Figure 3A-B; Supplementary Figure 3). Although *C. reinhardtii* has a relatively high concentration of lipids [17], only 188 (2.52%) of all CDS were associated with lipid metabolism. To understand how much genetic overlap there was among the five MAGs, we compared predicted genes identified with Prokka and clustered with OrthoVenn. The two *Limnohabitans* MAGs shared the highest number of genes (641), and across all MAGs 325 genes were shared (Figure 4). The *Pedobacter* MAG had the most unique genes in its gene set (117), while the Burkholderiaceae had so much overlap with other MAGs that it had relatively few unique genes (21).

**Figure 3.**
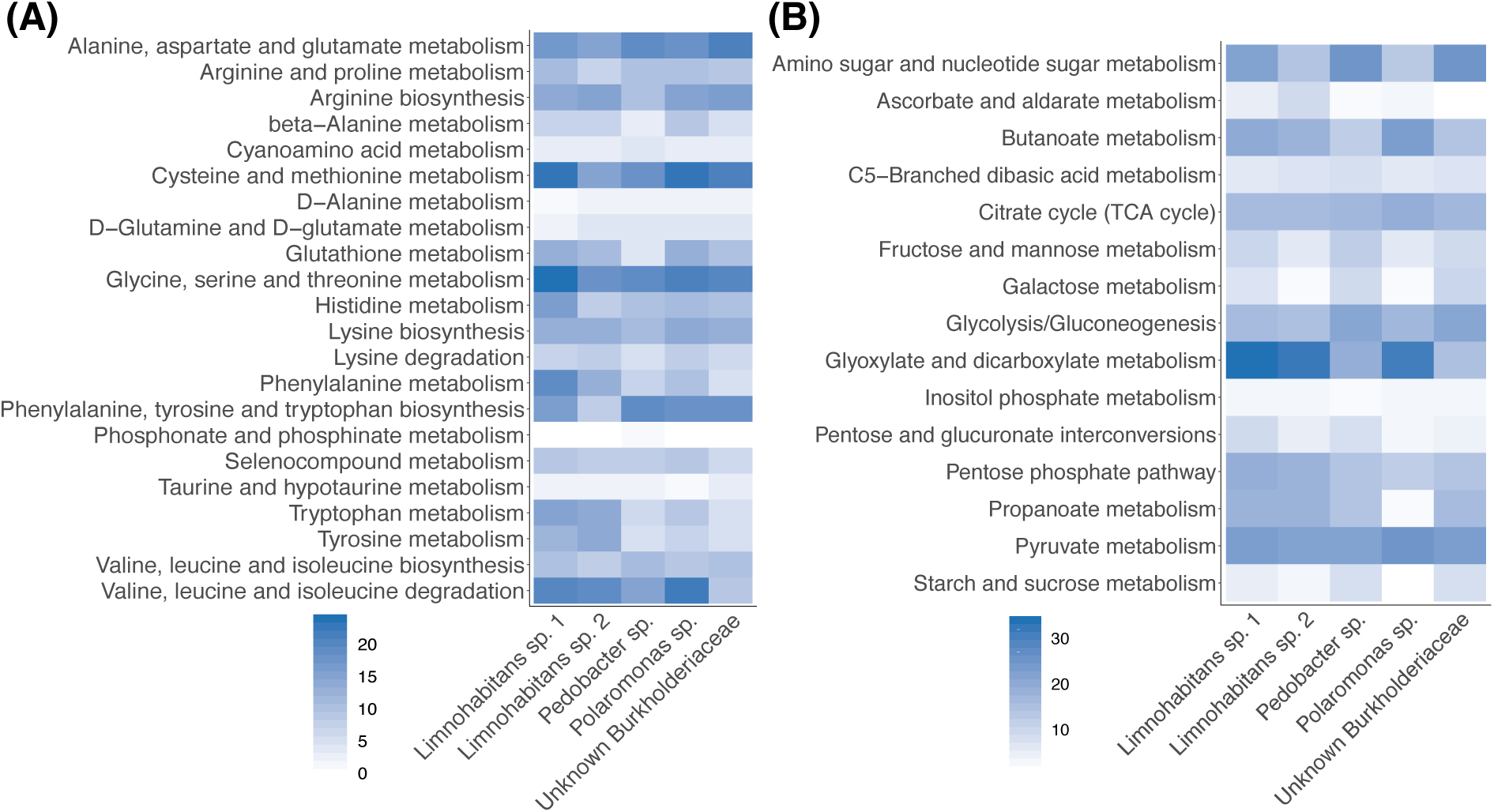
**(A)** Number of genes associated with carbohydrate metabolism pathways in the five MAGs. Genes within each MAG bin were annotated with Prokka, assigned KEGG orthology, and mapped to KEGG metabolic pathways. Genes assigned to pathways were then counted. **(B)** Number of genes associated with KEGG amino acid metabolism pathways in the five MAGs.

**Figure 4.**
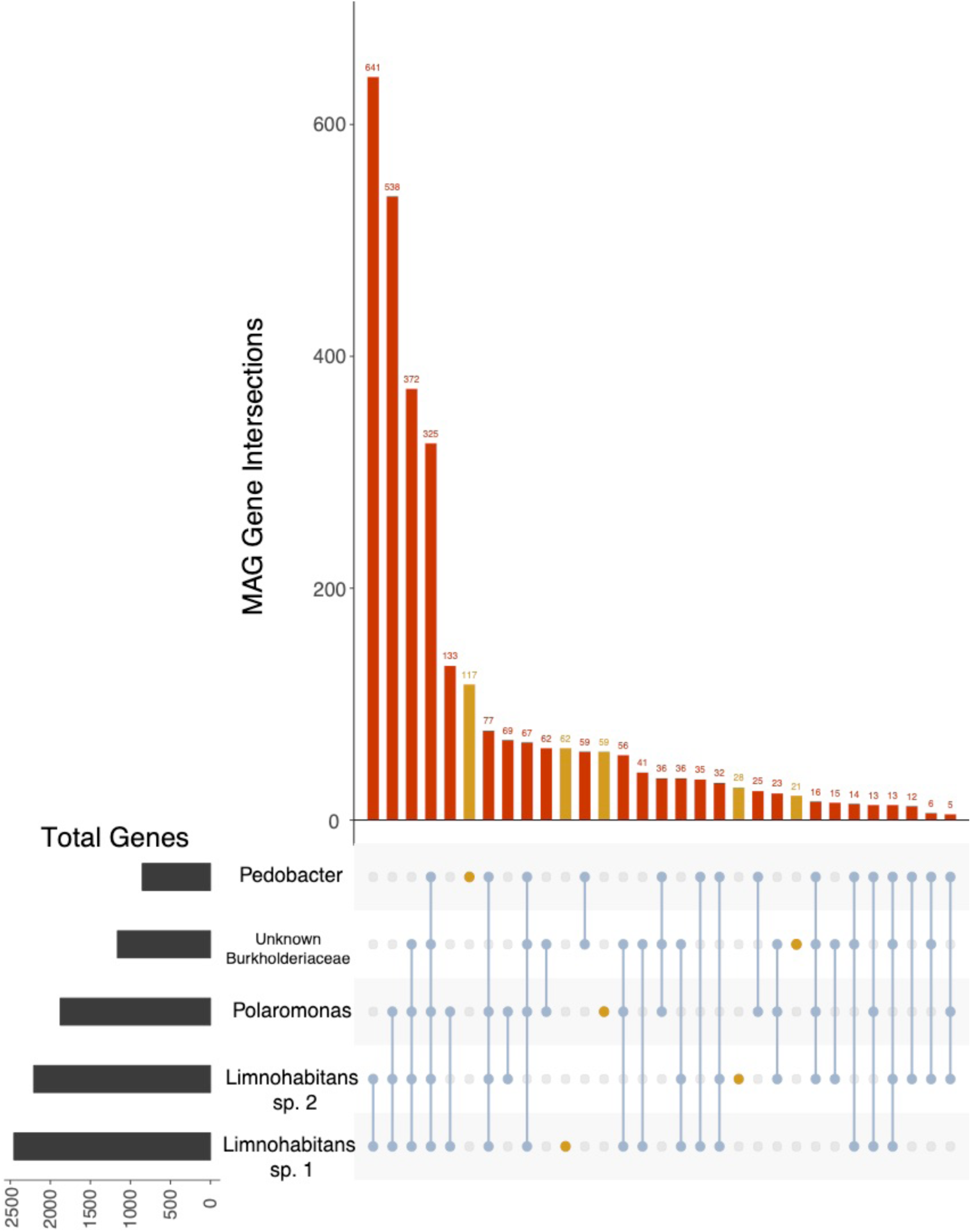
Shared and unique genes with KEGG orthologs in each MAG. Unique gene sets are highlighted in gold; shared genes are in red. Intersections between MAGs, or numbers of genes in each MAG’s gene set shared with the other MAGs connected by lines, are noted in the bottom matrix. The total number of genes in each MAG is listed on the left side of the matrix.

After examining genetic overlap and unique gene content of the five MAGs, we examined complete metabolic pathways annotated by KEGG present in the microbiota. Here, we describe some of the complete pathways annotated, specifically focusing on metabolic pathways involved in critical functions for bacteria such as nutrient uptake and biosynthesis as well as pathways that may be involved in host or environment interaction. A description and comparison of each MAG’s set of complete pathways can be found in Supplementary Materials and in Supplemental Tables 2-5. We separate complete pathways into those encoded by all or multiple MAGs, indicating shared pathways that could indicate functional redundancy or common metabolites accessible to multiple species, and pathways uniquely encoded by single MAGs, indicating potential niche differentiation within the microbiota [18]. Finally, we summarize some of the described shared and unique functions and some functions specifically associated with host association as genomic signals of host association, providing potential explanations for why these genera are commonly found in the *Daphnia magna* microbiota.

### Shared functions of the *Daphnia magna* microbiota

#### Nutrient uptake and major biosynthesis pathways

All five MAGs shared genes involved in some key metabolic pathways (Table 3, ‘All’ MAGs, Supplementary Tables 2 & 3). In all MAGs, a complete TCA cycle was encoded. Genes encoding a cytochrome *c* oxidase were identified in all MAGs, suggesting all have the capacity for aerobic respiration. All encoded for lipopolysaccharide transport and lipoprotein release. All MAGs except the *Pedobacter* encoded for a complete glyoxylate cycle, and the non-oxidative phase of the pentose phosphate pathway was present in all MAGs except the *Limnohabitans* species. Transporters for multiple TCA cycle intermediates are present across the MAGs, including a C_4_-dicarboxylate transport system in all MAGs that allows for transport of multiple different molecules into bacterial cells. Other transporters for TCA cycle intermediates encoded included those for alpha-glucosides, malate, fumarate, 2-oxogultarate, succinate, and aspartate.

**Table 3.**
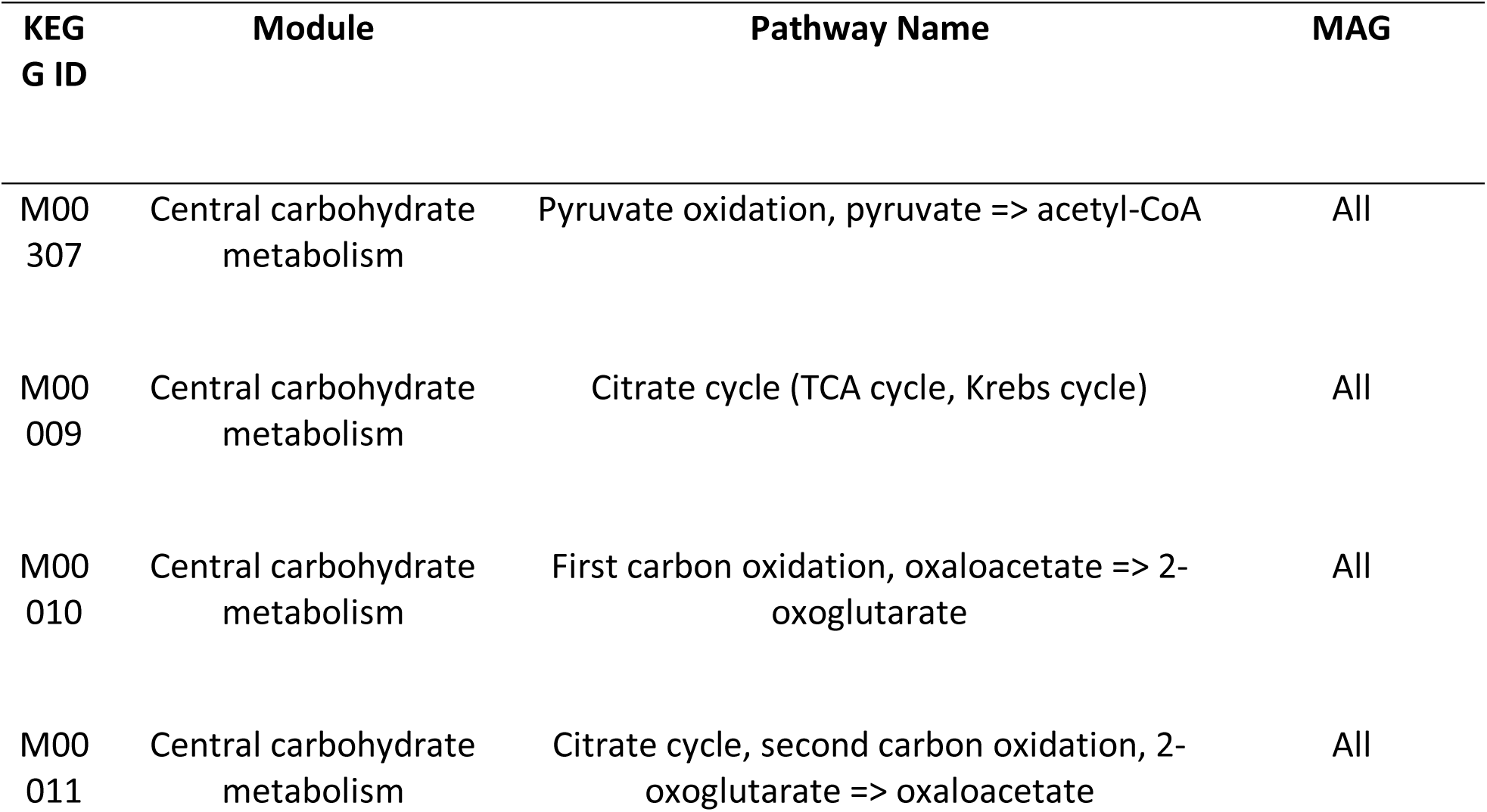
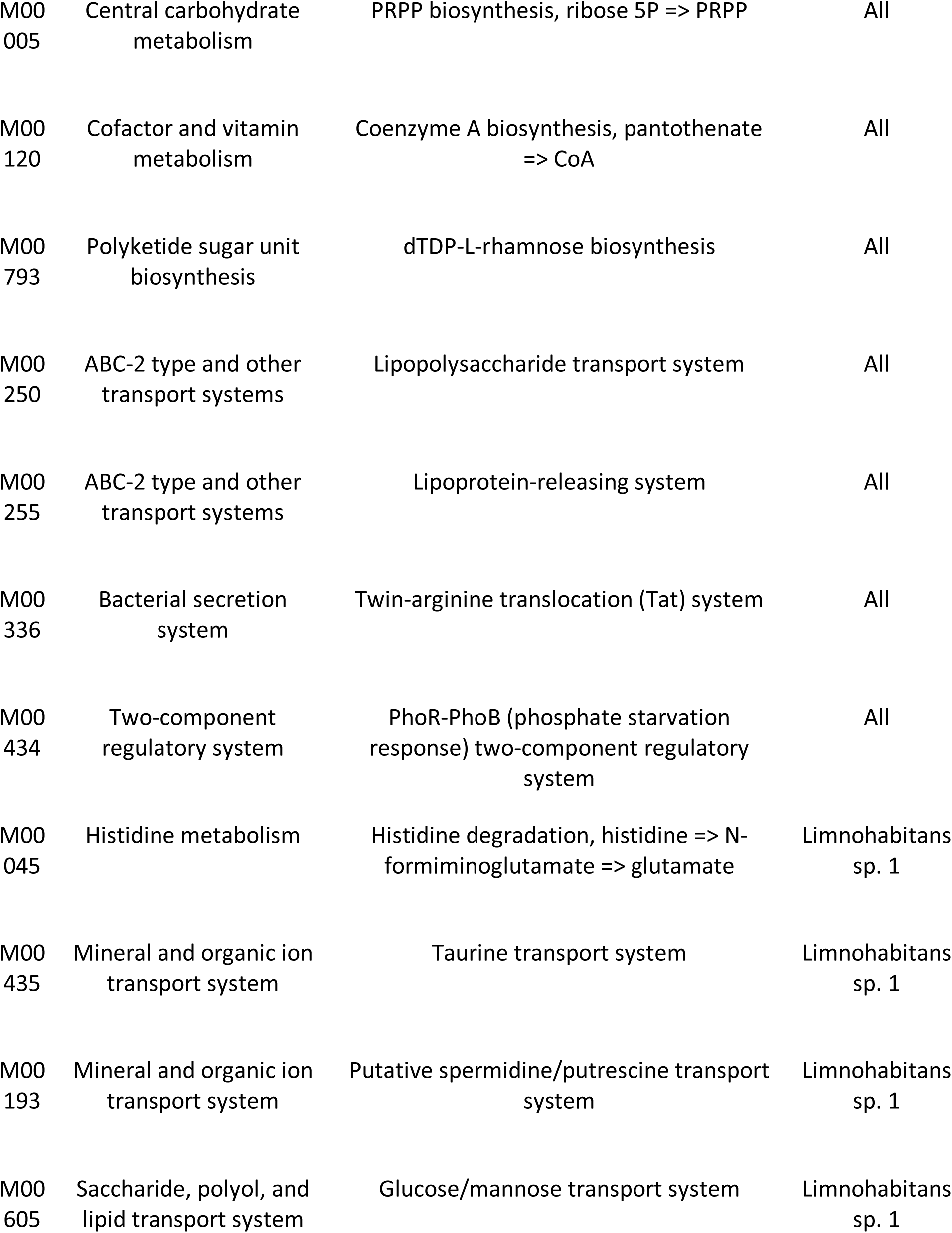
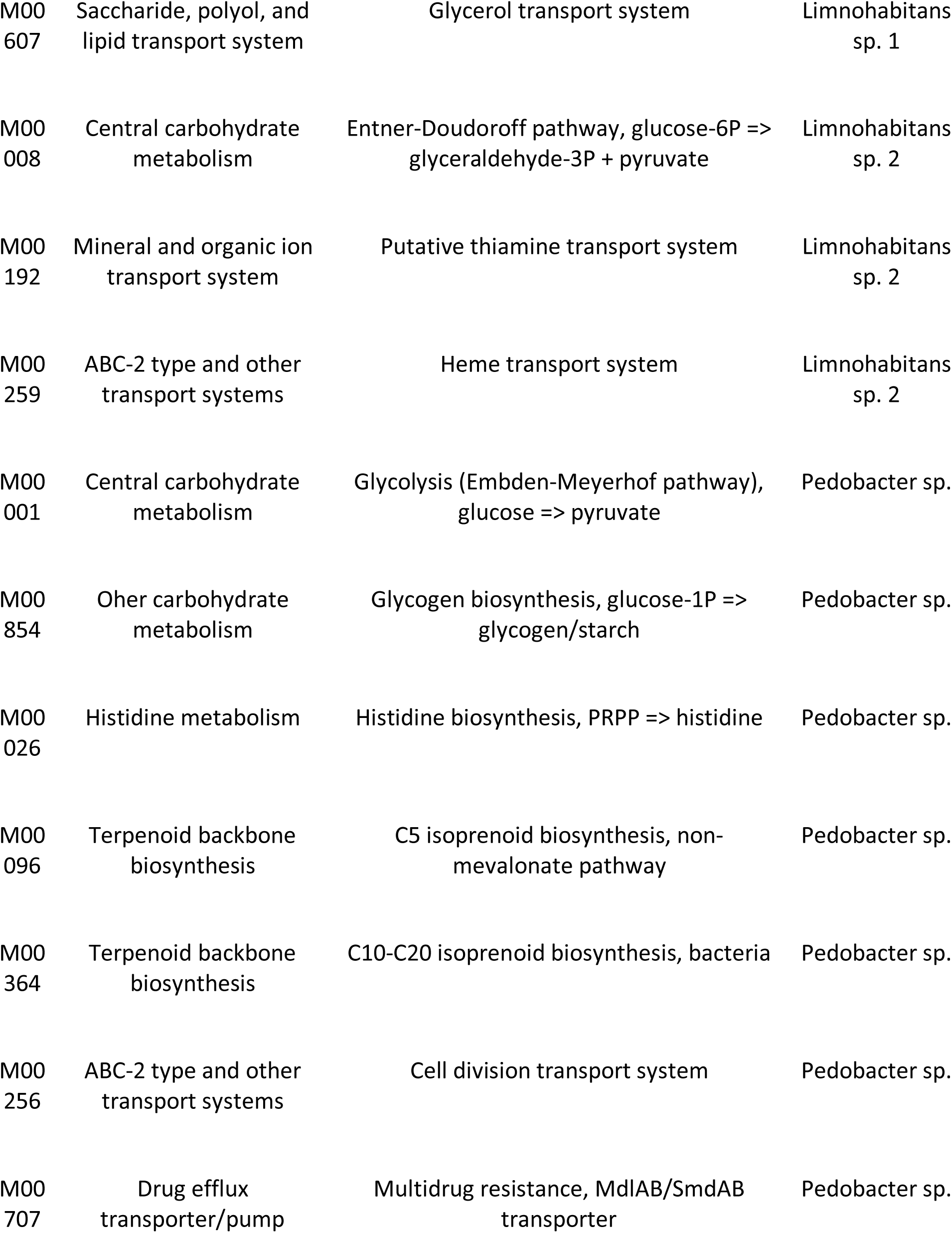
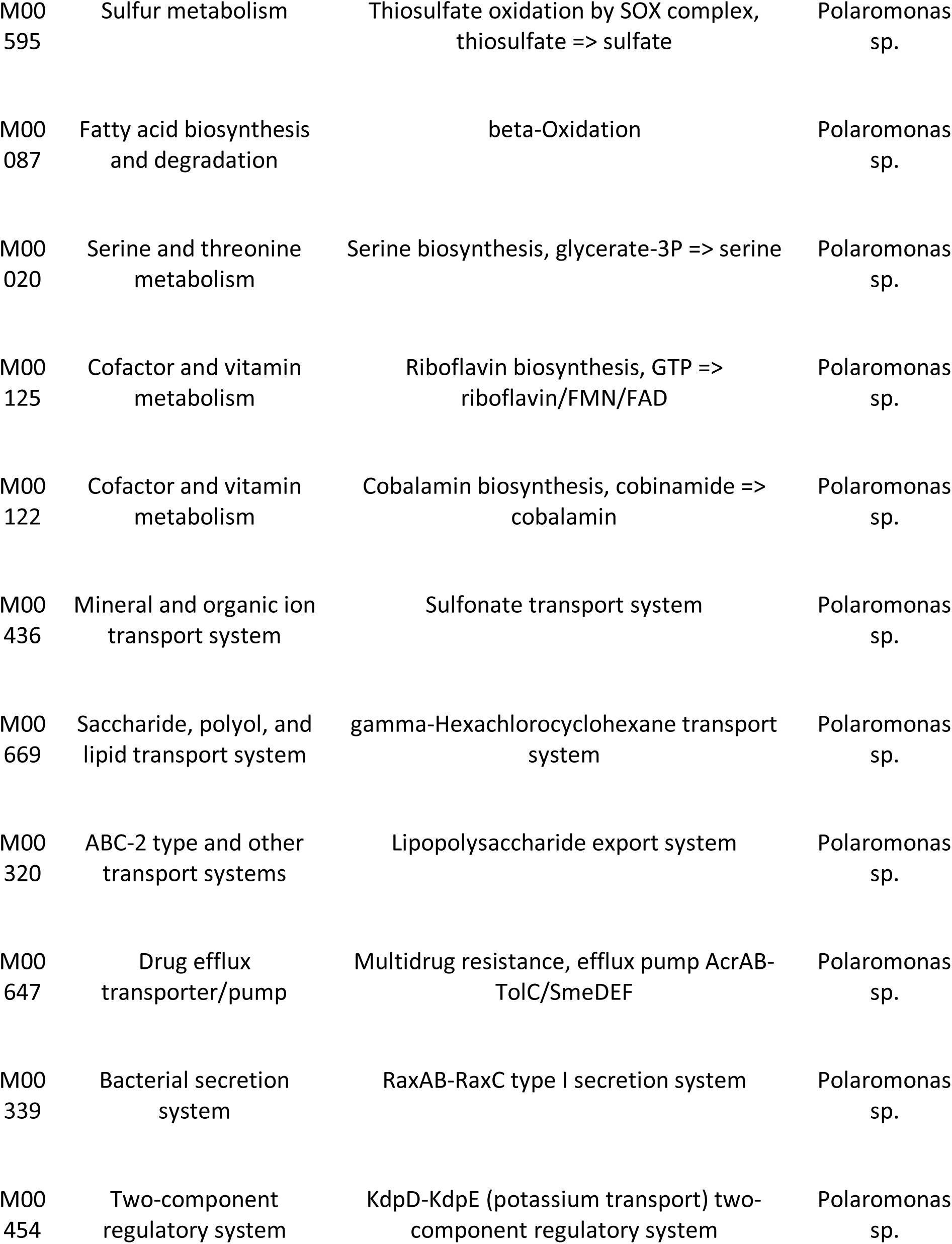
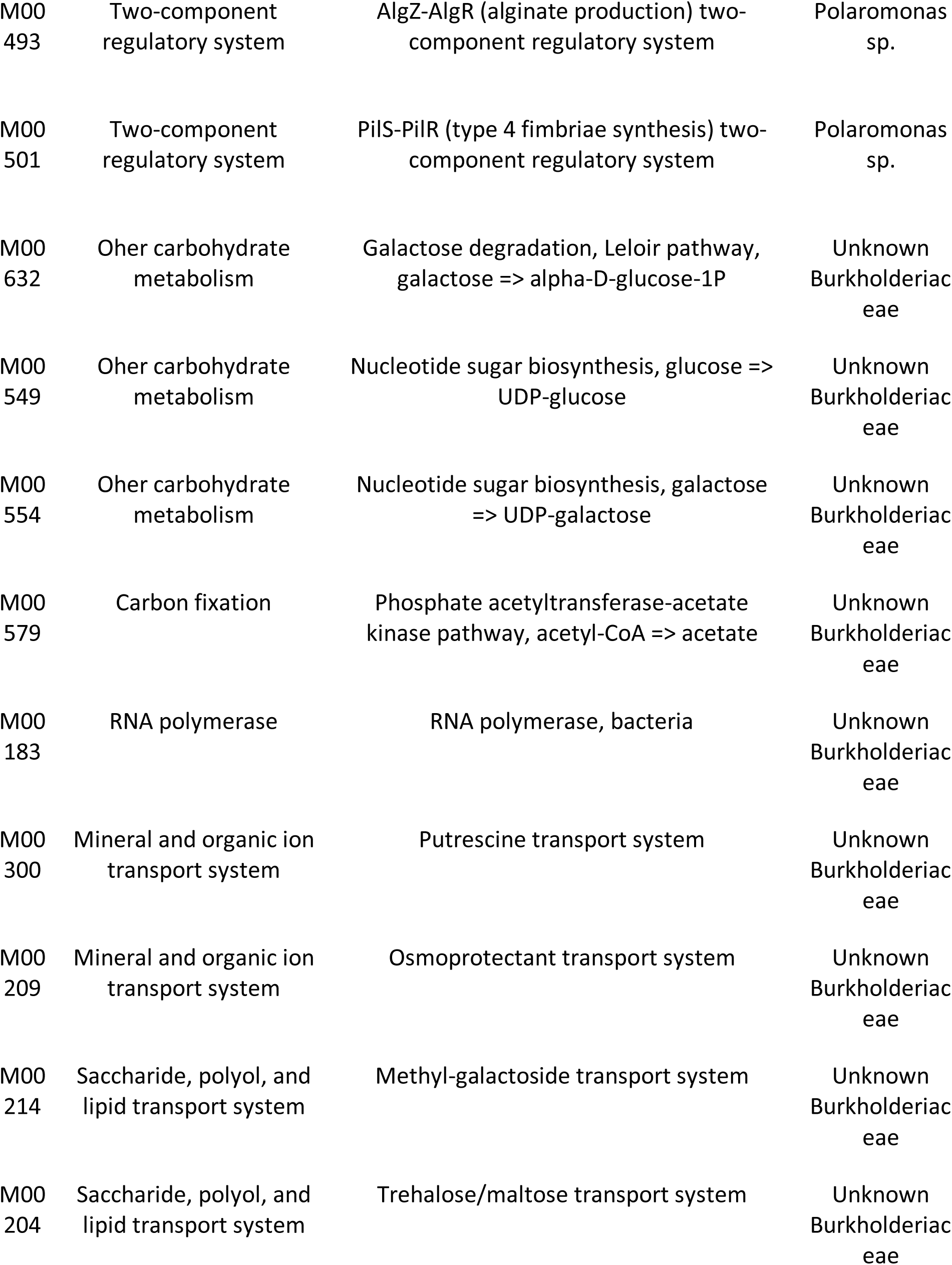
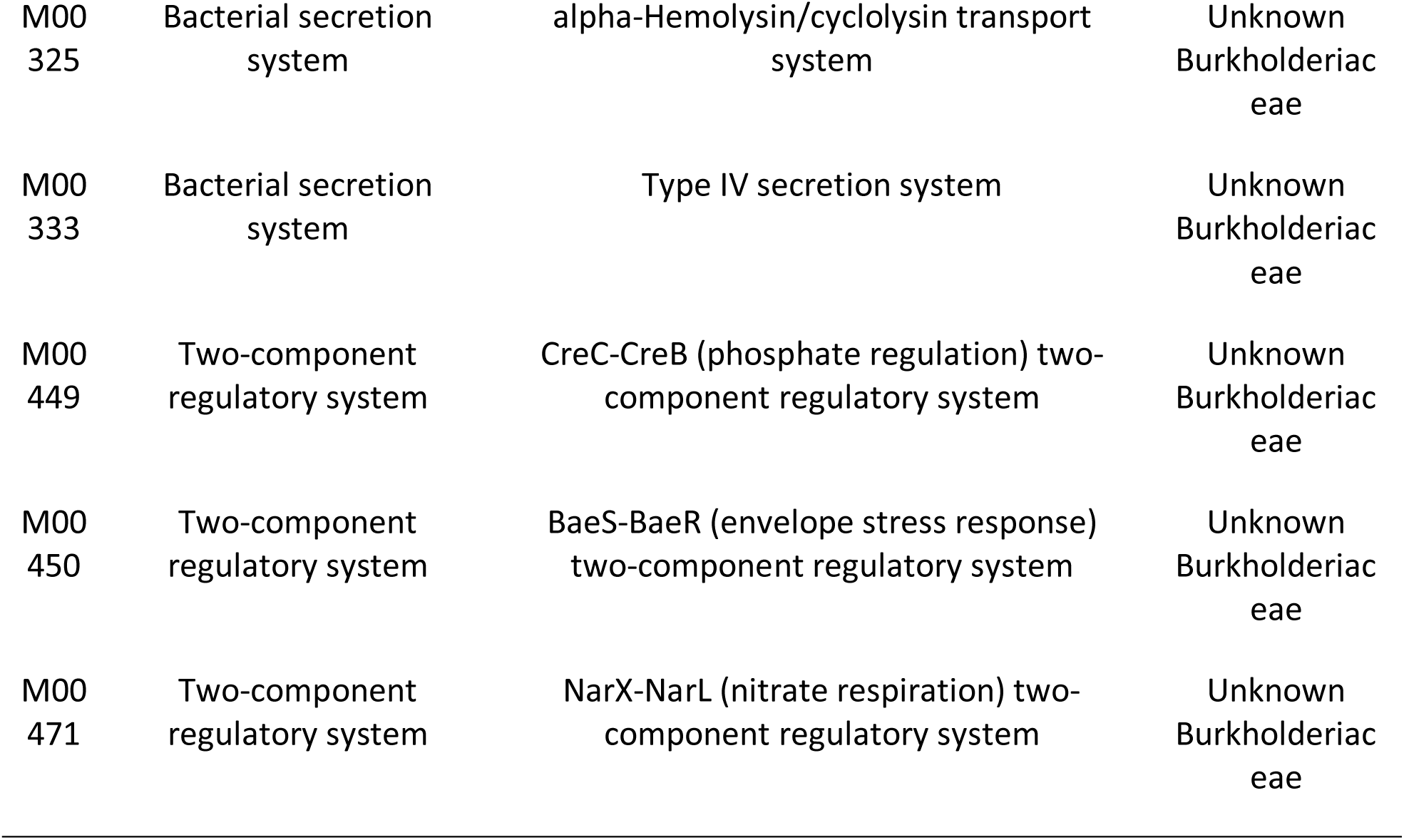
Shared and unique complete KEGG pathways in the five high- and medium-quality MAGs. ‘MAG’ column indicates which MAG encodes that complete pathway; ‘All’ denotes that the pathway is shared among all five MAGs.

Transport and biosynthesis of other essential molecules were shared across MAGs as well. All MAGs except *Pedobacter* shared transport systems for phospholipids, phosphate, and branched-chain amino acids. All species except for *Limnohabitans* MAG 2 encoded for the elongation step of fatty acid biosynthesis, but only the Burkholderiaceae, *Polaromonas*, and *Pedobacter* MAGs encoded for fatty acid biosynthesis initiation. The *Limnohabitans* MAGs shared multiple copies of sialic acid TRAP transporter genes (*siaM*, *siaT*, *siaQ*).

All genomes encoded for the transport of multiple necessary vitamins, including riboflavin (B_2_, *ribX*, *ribY*, *ribZ*), pantothenate precursors (B_5_, *panS*), and cobalamin (B_12_, *btuB*). Genes involved in the salvage pathway for cobalamin were also present in both *Limnohabitans* MAGs and in the *Polaromonas* MAGs (*cobO*, *cobP*). Some species were also able to biosynthesize vitamins: the *Pedobacter* and *Polaromonas* MAGs encoded for biotin biosynthesis, and all MAGs except *Pedobacter* encoded for tetrahydrofolate biosynthesis. All MAGs encoded for a pyridoxine 5-phosphate synthase (PdxJ), indicating likely biosynthesis of pyridoxine (B_6_).

Biosynthesis and transport of amino acids varied across MAGs, with no shared amino acid transport systems present across all species. All MAGs except *Pedobacter* were able to biosynthesize arginine and ornithine. The Burkholderiacae, *Limnohabitans* 1, and *Polaromonas* MAGs encoded for threonine, cysteine, lysine, proline, phenylalanine, tyrosine, and glutathione biosynthesis. Valine, leucine, and tryptophan were also able to be biosynthesized by some of the MAGs.

#### Host and environment interaction

Superoxide dismutase (*sodA*, *sodB*) and catalase-peroxidase (*katG*) were found in all MAGs. These act as reactive oxygen species detoxifiers and scavengers. The universal minimal Tat system was encoded by all MAGs (*tatA*, *tatC*), and both *Limnohabitans* MAGs also encoded *tatB*, allowing all of these species to transport folded proteins across their cell membranes. Complete adhesin transport systems and a gene associated with type IV pilus biosynthesis were present in both *Limnohabitans* MAGs and the *Polaromonas* MAG (*pilQ*). These MAGs also encoded two membrane proteases related to aminoglycoside resistance (HtpX, FtsH) [19]. All MAGs except the *Pedobacter* encoded for the QseC-QseB quorum sensing regulatory system. Though the *Polaromonas* MAG was the only one to encode for complete flagellar assembly, the *Limnohabitans* MAGs encoded for multiple genes involved in flagellar assembly (*flhA*, *flhB*, *flgB* – *L*). A gene for chemotaxis protein CheA was present in these three MAGs as well.

As none of the MAGs encoded for all necessary amino acid biosynthesis pathways, some amino acid importers were present across the MAGs. These include arginine (*artM*), proline (*proP*), cystine (*yecS*) in *Limnohabitans* MAGs, glutamine (*glnQ*, *glnM*) and glutathione (*gsiC*) in *Limnohabitans* MAGs and in the *Polaromonas* MAG, and histidine (*hisP*, *hisM*, *hisQ*) in the Burkholderiaceae MAG and *Limnohabitans* MAG 1. It is unclear whether food (e.g., *Chlamydomonas reinhardtii*) or export from other bacteria is the source of these amino acids for these MAGs. As some MAGs are able to biosynthesize amino acids that others must import, there may be cross-feeding among species in the microbiota occurring.

Environmental stress tolerance mechanisms were shared among genomes as well. All encoded for the phosphate starvation response regulatory system PhoR-PhoB. The *Limnohabitans* MAGs and the *Polaromonas* MAG shared the EnvZ-OmpR osmotic stress response system, allowing the bacterial cells to respond to changes in osmolality. *Limnohabitans* MAG 2 and the *Polaromonas* MAG shared CusS-CusR, a copper tolerance regulatory system.

### MAG-specific functions of the *Daphnia magna* microbiota

#### Nutrient uptake and major energy pathways

There were potential differences in respiration for some of the MAGs. The Burkholderiaceae encoded a nitrate/nitrite transporter (*narK*) and a respiratory nitrate reductase (*narGHI*). Use of this system for respiration was supported by the presence of genes encoding the nitrate respiration two-component regulatory system NarX-NarL. The *Polaromonas* encoded for thiosulfate transport (*cysT*, *cysW*, *cysP*) and potential use via a subunit of the Sox complex (*soxA*). We found a supplementary energy pathway in the *Limnohabitans* MAGs, which share a pathway for the biosynthesis of bacteriochlorophyll (*pufLM*, *bchY*). This pathway has been documented in other *Limnohabitans* species [20].

Differences in carbohydrate metabolism among MAGs was apparent. The *Pedobacter* MAG was the only high-quality genome to encode for glycolysis and glycogen biosynthesis; the Burkholderiaceae MAG for the Leloir pathway; and *Limnohabitans* MAG 2 for the Entner-Doudoroff pathway. The *Polaromonas* MAG encoded for a complete beta-oxidation system, enabling fatty acid metabolism. Different carbohydrate transporters were found in the MAGs as well. *Limnohabitans* MAG 1 uniquely encoded for the transport of glucose, mannose, and glycerol (*gtsABC*, *malK*).

It also encoded for transport of L-arabinose (*araPQ*) and for a semiSWEET general sugar transporter. The Burkholderiaceae MAG encoded for methyl-galactoside transport (*mglABC*) and a galactose processing pathway. The *Pedobacter* MAG can potentially utilize chitin, as the MAG uniquely encoded for chitin degradation to fructose-6-phosphate (*chiA*, *chb*, *nagB*).

Other pathways for vitamin and amino acid import, biosynthesis, and degradation were uniquely present in some genomes. *Limnohabitans* MAG 1 was the only MAG to encode taurine import (*tauA*, *tauB*). This MAG also uniquely encoded a pathway for the degradation of histidine to glutamate. *Limnohabitans* MAG 2 could transport thiamine into the cell via a putative thiamine transport system (KEGG Module M00192). Uniquely, *Polaromonas* could synthesize cobalamin (*cobA*, *cobQ*, *cbiB*, *cobP, cobC*). It may also synthesize riboflavin via the purine biosynthesis pathway.

#### Host and environment interaction

Multiple genes and pathways involved in antibiotic resistance and detoxification were found in the bacterial MAGs. *Pedobacter* encoded an MdlAB/SmdAB transporter as well as two genes involved in macrolide (*macA*, *macB*). The *Polaromonas* MAG encoded an AcrAB-TolC/SmeDEF efflux pump, and the Burkholderiaceae MAG encoded the BaeS-BaeR envelope stress response two-component regulatory system.

The MAGs encoded for different secretion systems and regulatory systems. The *Polaromonas* MAG was the only one to encode for a type I secretion system (RaxAB-RaxC). The Burkholderiaceae MAG encoded for an alpha-hemolysin and cyclolysin secretion system and a type IV secretion system (*virB1* – *5*, *10*, *11*). The Burkholderiaceae MAG encoded for an osmotically-inducible protein, OsmY, part of an osmoprotectant ABC transporter complex, and a putrescine transporter (*potFGHI*).

### Genomic signs of host association in the *Daphnia magna* microbiota

Because the *Daphnia magna* microbiota has been shown to differ significantly from the surrounding aquatic environment at the genus level [9,21], we examined the MAGs for any potential indicators of host association. Multiple instances of host immune system evasion or tolerance were noted in the MAGs. Both *Limnohabitans* MAGs encoded for the microbial stealth protein CpsY, implicated in host immune system evasion [22]. Genes involved in dTDP-L-rhamnose biosynthesis were present in all MAGs, indicating these species are able to use rhamnose as an alternative cell wall polysaccharide. L-rhamnose has been linked to bacterial viability and virulence [23], and similarly structured polysaccharides have been shown to modulate host immune systems [24]. A gene involved in quorum quenching (signaling between other microbial species or between microbes and the host) was identified in *Limnohabitans* MAG 1 and the *Pedobacter* MAG (*ytnP*). Other virulence factors and regulators were identified in both *Limnohabitans* and the *Pedobacter* MAG (*cvfB*, *bvgS/bvgA*).

These bacterial species may benefit the host via amino acid and vitamin biosynthesis and export. Complex eukaryotes must acquire some essential vitamins and amino acids from their diet or from heterotrophic microorganisms. Six amino acids have been demonstrated as essential for *Daphnia magna’s* close relative, *Daphnia pulex*: arginine, histidine, leucine, phenylalanine, isoleucine, and tryptophan [25]. All of the dietary essential amino acids are biosynthesized by at least one of the MAGs, and some may be able to export these as well. The *Limnohabitans* MAGs and the Burkholderiaceae MAG encode for an arginine exporter (*argO*), *Limnohabitans* sp. 1 encodes for a threonine exporter (*rhtA*), and the Burkholderiaceae MAG a threonine-serine exchanger (*steT)*. The *Chlamydomonas reinhardtii* food source used here does not contain cobalamin [26], a beneficial vitamin for *Daphnia magna*, suggesting that *Daphnia magna* raised in culture must acquire cobalamin from the culture media and from microorganisms. In the *Daphnia* metagenome, the *Polaromonas* MAG encodes for cobalamin biosynthesis. Other vitamins are biosynthesized by at least one of the MAGs, including tetrahydrofolate, biotin, and pyridoxine. Supplementation of *Daphnia magna* growth media with biotin and cobalamin has been shown to increase host fitness [27], though it is unclear if these vitamins are essential for *Daphnia* survival.

Other host-microbe interactions were also present in the MAGs. An essential nutrient for bacteria is iron, and catalase genes were found in all MAGs along with other ABC transporters for iron. The *Limnohabitans* MAG 2 encoded for heme binding proteins and a heme export system (*ccmB*, *ccmC*), systems implicated in bacterial use of host-synthesized heme [28]. The *Pedobacter* MAG encoded for an N-acetylneuraminate lyase (*nanA*) and all five MAGs contain genes involved in transport of sialic acids. As sialic acids are found in complex host tissue [29], this may indicate cleavage of sialic acids from host cells for import and use by the bacteria. Pathways involved in host invasion and colonization were also present in the MAGs. The *Polaromonas* MAG encoded a suite of type IV pilus and fimbriae associated genes, including type IV pilus biogenesis factors (*pilY1*, *pilQ*), fimbrial proteins (*pilE*), the type IV fimbriae synthesis two-component regulatory system PilS-PilR, and a complete set of flagellar assembly genes. This MAG, along with the two *Limnohabitans* Mags, also encoded for twitching motility (*pilT*). Both *Limnohabitans* MAGs encoded for swarming motility (*swrC*, *rssA*), and all three for other genes involved in chemotaxis (*pctA*, *cheA*, *cheY*, *tar*, *cheB*, *mcp4*, *tsr*, *cheW*). Though genes for adhesin production were not identified, genes involved in adhesin transport were identified in both *Limnohabitans* MAGs and in the Burkholderiaceae MAG (*bmaC*, *ehaG*, *ata*).

## Discussion

To date, studies examining the *Daphnia magna* microbiota have only sequenced the 16S rRNA marker gene to understand broad-level interactions and functions of the microbiota. Here, we were able to assemble the first metagenome-assembled genomes from the *Daphnia magna* microbiota to elucidate genome-specific functions associated with highly abundant members of the bacterial community. Our 16S rRNA gene level data shows that the *Daphnia magna* bacterial community is structured differently than the surrounding environment and from the microbiome of their food, which has been also been corroborated by earlier studies [9,30]. We identified five MAGs that were distinct from each other and distinct from their closest sequenced relatives based on average nucleotide identity. We found that these MAGs were high- or medium-quality, meaning they contained some or most of the single-copy genes found in all bacteria. With metagenomic short-read shotgun sequencing it is unlikely that the MAGs assembled here contained the full gene sets associated with these species; however, bins of relatively high-quality suggest that we found many important genes within each assembled genome. Most studies of the *Daphnia magna* microbiota using marker-based sequencing have identified more OTUs or ASVs and higher diversity than the 12 bins we identified, likely due to the higher sequencing depth necessary for shotgun sequencing to identify rare taxa [31]. However, the five MAGs assembled from our shotgun sequencing are mostly consistent with genera found in 16S rRNA sequencing from other studies and from our own sequencing.

The Burkholderiales order has been demonstrated as the most abundant in the *Daphnia* gut and whole organism microbiota [9,13], and four of the five MAGs assembled here were identified to families or genera within this order. Two *Limnohabitans* MAGs were assembled and found at high abundances in adults and juveniles. Within the Burkholderiales, the genus *Limnohabitans* has been reported to be highly abundant in the *Daphnia* microbiota, and has been implicated in increasing host fecundity [12,15]. Here, we show that there are two distinct *Limnohabitans* MAGs that encode for different metabolic potential. Only one other study has definitively identified more than one OTU in this genus [13]. We also identify two other MAGs in the Burkholderiales order, including a *Polaromonas* species and one unclassified Burkholderiaceae. Surprisingly, we also identify a *Pedobacter* MAG, which has previously only been reported as a rare taxon in the *Daphnia* microbiota [32].

Analysis of annotated genes and pathways across the five MAGs showed overlap and differences in metabolism. Key pathways such as the TCA cycle were shared across all MAGs, and multiple ABC transporters of key vitamins and amino acids were identified. Many genes encoding for the use of different carbohydrates were encoded across the MAGs. Microalgae are generally rich in carbohydrates [33] and serve as a major food source for *Daphnia magna* [34], potentially allowing these microbes easy access to myriad nutrient sources. The two *Limnohabitans* MAGs shared a high proportion of annotated genes, suggesting some functional similarity between them. Indeed, multiple metabolic pathways were shared between these two MAGs, including the glyoxylate cycle, bacteriochlorophyll biosynthesis, and biosynthesis of some vitamins and amino acids. The *Limnohabitans* MAGs also shared many genomic features with the *Polaromonas* MAG and the Burkholderiaceae MAG, including multiple TCA cycle intermediate importers and several transport systems. There is also potential flexibility among the taxa in encoded respiration, notably in the *Polaromonas* MAG’s ability to use thiosulfate and the Burkholderiaceae’s ability to utilize nitrate under hypoxic conditions. Along with the variety of two-component regulatory systems found within each MAG, the wide range of potential respiratory pathways may allow functions to sustain through different bacterial species even when stressful environments cause fluctuations in the abundance of species within the microbiota.

The difference between taxa identified in the *Daphnia magna* microbiota and its environment suggest that there are some key interactions between the host and its associated microbes in order to establish and maintain these microbial populations. Furthermore, there may be some interactions between bacterial species in the microbiota that could impact the host. We found many genes in the *Limnohabitans* MAGs and the *Polaromonas* MAG involved in flagellar assembly, type IV pilus biogenesis and production, and biofilm formation, all of which have been implicated in host colonization and successful adhesion to host-associated surfaces [35,36,37]. All MAGs encode for l-rhamnose production, which has been implicated in adhesion to other cells [23]. Genes in secretion systems implicated in host cell adhesion, particularly Type I, II, and IV were also encoded [38]. We found many genes involved in host immune system evasion or modification, which may allow these bacteria to persist within the host species [39]. Notably, superoxide dismutase and catalase were encoded by multiple MAGs, suggesting the bacteria could defend against radical oxygen species produced by the host as a defense mechanism [40]. Also present across MAGs are genes involved in the detoxification of antibiotics and toxins, including multidrug efflux transporters and pumps (MdlAB/SmdAB, AcrAB-TolC/SmeDEF), macrolide export (*macA*, *macB*), and stress tolerance to antibiotics (BaeS-BaeR).

Biosynthesis and provision of amino acids by bacteria to their host is a well-documented set of interactions that is known to confer fitness benefits to the host [41–43]. Here, we find that the *Limnohabitans* MAGs and the Burkholderiaceae MAG encode for the export of arginine and threonine, essential amino acids for the host *Daphnia* [25]. Similarly, biosynthesis or metabolism of vitamins and minerals by bacteria and provisioning to the host has been well-documented in the microbiota of other organisms [44–46]. Many genes involved in vitamin B biosynthesis were found across all MAGs. Media supplemented with cobalamin (B_12_) is used to successfully culture *Daphnia magna* [47], and we find the *Polaromonas* MAG encoded for a complete cobalamin biosynthesis pathway, suggesting potential vitamin provisioning to the host from this species. We also find a potential microbe-microbe interaction, where the *Pedobacter* MAG encodes for cleavage of sialic acids from host tissue, where it can then be transported and utilized by other species as a carbohydrate source via a sialic acid TRAP transporter. The breakdown of sialic acid to metabolites that are accessible to the host by microbiome-associated species has been shown to increase host fitness [48]. If *Limnohabitans* are able to use sialic acid as a nutrient source, this may be the basis for a microbe-host-microbe interaction, where *Limnohabitans* provides essential amino acids to the host using energy generated from metabolism of molecules provisioned from the host.

In total, our data shows that there is much versatility in metabolism among the MAGs, but some overlap in function. As *Daphnia magna* are indiscriminate filter feeders, they may feed on a wide variety of particulate matter with variable nutrient profiles. The versatility in metabolism encoded by these MAGs indicates that they are able to utilize this unpredictable range of nutrients both in the digestive tract and on the carapace. The specific functions of certain MAGs, particularly in amino acid and vitamin biosynthesis and export, seem critical in providing nutritional benefits to the host zooplankton.

## Conclusions

*Daphnia magna* is an important model system for multiple facets of ecology, and has recently become an organism of interest for understanding fundamental questions about the microbiota. Our metagenomic sequencing and subsequent analysis characterizes the *Daphnia magna* microbiota to the species level and finds some genomic features that allow core bacterial species to acquire and biosynthesize nutrients, and to potentially interact with their host via amino acid and vitamin export. By examining this relatively simple microbiota via metagenome-assembled genomes, we can begin to investigate metabolic interactions between the host and its associated microbes. Future work to further elucidate functions of these MAGs will involve long-read metagenome sequencing to complete genome assemblies and pure, single-isolate sequencing to understand strain variation within the microbiota. Furthermore, transcriptomics and metabolomics could be used to understand which of these encoded genes are functioning under different environmental and host conditions, and will direct future hypotheses on host-microbiota interactions. For example, how much of the differences in *Daphnia* life history and population dynamics across food environments [49] can be attributed to differences in microbiota composition? These results will help to inform future work studying the effects of the microbiota on host health and population dynamics across ecological contexts. Moreover, as more populations of *Daphnia* and their microbiota are sequenced, it will become possible to examine the coevolutionary relationships between hosts and microbiota, and this functional information will be essential for making sense of those relationships.

## Methods

### Sample collection and extraction

Two samples of 100 21-day-old adult *Daphnia magna* and two samples of 6-day-old juvenile Daphnia were collected from laboratory cultures maintained in defined COMBO medium [47]. Laboratory cultures were fed a standardized volume of green algae *Chlamydomonas reinhardtii* (CPCC 243) to provide 0.25 mg C/ml/day. Samples were immediately ground after collection using in sterile 1.5mL microcentrifuge tubes. To separate bacterial cells from host cells, a modified protocol from Benson et al. 2014 was used [50]. Samples were suspended in 2mL PBE buffer and layered on a cushion of 50% sucrose, then centrifuged at 4,000g for 10 minutes. DNA was extracted from the pellets at the base of the sucrose fractions following the Qiagen DNEasy Blood & Tissue Kit spin-column protocol of total DNA from animal tissues (Qiagen, Hilden, Germany).

### Library preparation and sequencing

Shotgun sequencing libraries from the two adult samples and two technical replicates of one juvenile sample were prepared and multiplexed using the Illumina Nextera XT kit and protocol. Input DNA was quantified using the Qubit dsDNA system. Libraries were checked using the Agilent Technology 2100 Bioanalyzer. Libraries were manually normalized due to a final library yield under 10nM. Paired-end sequencing was performed on an Illumina MiSeq using a MiSeq Reagent Kit V2.

### Quality filtering and metagenome assembly

Reads were demultiplexed using the built-in Illumina MiSeq Reporter. Quality of demultiplexed reads was checked using FastQC v0.11.5. Reads were trimmed using Trimmomatic v0.36 [51] with the commands: ILLUMINACLIP:NexteraPE-PE.fa:2:30:10 LEADING:3 TRAILING:3 SLIDINGWINDOW:4:25 MINLEN:36 to remove adapter sequences and to remove segments of reads where quality fell below 25. Trimmed reads were mapped to the Daphnia magna draft genome [52] using BWA [53], Samtools [54], and BEDtools [55], and reads with greater than 80% identity over 50% of the read were filtered. Remaining paired unmapped reads were assembled de novo using metaSPAdes in SPAdes v3.11 [56]. A co-assembly of all four samples, a co-assembly of the adult samples, a co-assembly of the juvenile samples, and individual assemblies for each sample were created.

### Taxonomy-independent sequence binning and identification of individual metagenome-assembled genomes

To resolve genomes of identified organisms and to discover genomes of organisms not present in read-based taxonomy identification programs or entirely new organisms, contigs from the master co-assembly with reads mapped from each sample were binned using CONCOCT within Anvi’o [57], accepting contigs over 2500 bp in length. Contigs greater than 20kb in length were split into 20kb fragments prior to running CONCOCT. Bins from the master co-assembly were assessed using Anvi’o, using Parks et al.’s quality score (genome completeness - 5x estimated redundancy or contamination) of ≥50 as a cutoff for further refinement [58]. Bins meeting this quality cutoff were manually curated within Anvi’o, where contigs within a bin that deviated dramatically from the mean GC content or mean coverage of the bin were removed from the bin. Bins that increased completeness when merged, did not increase redundancy above 10%, and were similar in GC content were merged. Bins that did not have high (>90% completion) or medium quality (>50% completion) after merging and refining were not analyzed further. After merging and refining, bins that still met the quality score cutoff were assigned taxonomy using GTDB-Tk [59,60]. GTDB-Tk uses average nucleotide identity and genome topology to find the closest genomic relative in its database. The same process was repeated for the juvenile co-assembly and the adult co-assembly to confirm species presence and attempt to resolve species identity at different host life stages. Similarity between MAGs was calculated using the average nucleotide identity tool in Pyani [61].

### Read-based taxonomic classification

Kaiju v1.5 [62], Kraken v1.0 [63], and MetaPhlAn2 v2.6 [64] were used to assign taxonomy to reads. We used all three identifiers due to due to Kaiju’s high rate of false identification [65] and MetaPhlAn2’s use of specific marker genes from reference organisms rather than entire genomes. All programs were used with their built-in databases. Kraken results were confirmed via cross-comparison of abundant species with both Kaiju and MetaPhlAn2. All three taxonomy profilers were also used to assign taxonomy to contigs assembled from each sample. Visualization of each sample’s community composition was performed in R.

### Functional profiling

Contigs from each metagenome-assembled genome and from the adult, juvenile, and master co-assemblies were annotated using Prokka v1.12 [66]. Genes annotated in Prokka were assigned KOs (KEGG Orthologs) using the GhostKOALA tool on the Kyoto Encyclopedia of Genes and Genomes [67]. KOs identified from GhostKOALA were mapped to standard KEGG categories and metabolic pathways using the KEGG Pathway Mapper & KEGG Module tools to examine pathway completeness and identify pathways of interest [68]. Genes identified using KEGG and GhostKOALA in each MAG were compared using OrthoVenn [69]. Overlapping and unique orthologs were compared using custom R scripts and with the ‘UpSetR’ package [70].

### 16S rRNA gene sequencing and identification of contaminant taxa

The V4 hypervariable region of the 16S rRNA gene was sequenced on the Illumina MiSeq using a MiSeq Reagent Kit V2 and the same Qiagen DNEasy Blood & Tissue Kit and reagents as in the shotgun sequencing sample processing. Four samples of five adult *Daphnia magna* were sequenced to compare community composition found in 16S sequencing to that found in shotgun sequencing, along with four samples of the COMBO media *Daphnia magna* cultures are raised in, four samples of *Chlamydomonas reinhardtii*, and two samples of the DNA sequencing kit and library preparation kit as negative controls. Paired-end reads were analyzed in R using the ‘dada2’ package to trim primer sequences, identify amplicon sequence variants, and assign taxonomy [71]. Taxonomy was assigned using the RefSeq+RDP taxonomic training data set formatted for dada2 [72]. Further analysis of community composition and visualization were carried out using the ‘phyloseq’ package in R [73].

## Supporting information

Supplemental Descriptions of MAGs, Supplemental figures, and tables

## Declarations

### Ethics approval and consent to participate

Not applicable.

### Consent for publication

Not applicable.

### Availability of data and materials

The datasets supporting the conclusions of this article are available at the NCBI BioProject Portal under IDs PRJNA543317 (shotgun sequences) and PRJNA543842 (16S rRNA sequences. Raw metagenomic sequencing reads from each sample are deposited under accession numbers SAMN11660785 and SAMN11660786. 16S rRNA sequencing data can be found under accession numbers SAMN11784745 - SAMN11784837. All scripts for data analysis and visualization are available on GitHub (https://github.com/reillyowencooper/daphnia_magna_metagenome).

### Authors’ contributions

ROC and CEC designed the study. ROC collected and analyzed the metagenomic data. ROC wrote the first draft of the manuscript, and ROC and CEC revised the manuscript and approved its final form.

## Acknowledgements

We are grateful for Dr. Andrew K. Benson, Mallory Van Haute, and Qinnan Yang’s assistance with sequencing preparation and machine use, and to Brady Bathke for assistance with bioinformatic analyses. This work was completed utilizing the Holland Computing Center of the University of Nebraska, which receives support from the Nebraska Research Initiative.

