## Supplemental Descriptions of MAGs, Supplemental figures, and tables for "Characterization of core bacterial species in the *Daphnia magna* microbiota using shotgun metagenomics"

**MAG profiles**

*Limnohabitans* sp.1

*Nutrient uptake and major energy pathways*

*Limnohabitans* sp.1 is able to utilize a wide range of molecules for nutrients, similar to many other *Limnohabitans* species (CITE limno paper with 'metabolic potential'), and is likely to rely mainly on short-chain fatty acids, carboxylates, and amino acids for energy. It carries transporters for glucose, mannose, glycerol, and alpha-glucosides, but does not seem to be able to catabolize these as there are no complete carbohydrate-processing pathways encoded in the genome. A phosphofructokinase, phosphoglycerate kinase, and enolase are not present in the *Limnohabitans* MAG, indicating that glycolysis is likely not functional. The pentose phosphate pathway and the Entner-Doudoroff pathway, two alternative pathways for energy production via carbohydrates, are not complete either.

While no complete carbohydrate-processing pathways are present, it is possible that the genes were simply not captured by metagenomic assembly. *Limnohabitans* may make up for the lack of full carbohydrate-processing pathways through production of bacteriochlorophyll *a*, which supplements bacterial metabolism with carbon acquired from light energy (Yurkov, 2009). This has been demonstrated in other species of *Limnohabitans* (Kasalicky, YEAR), but not in a host-associated species. Bacteriochlorophyll utilization by this species may fix carbon for energy, which in turn may lower demand on the host for other nutrients. More generally, *Limnohabitans* sp. 1 contains cytochrome *c* oxidase genes, indicating aerobic metabolism.

Though it is not apparent that this species is capable of carbohydrate catabolism, it does appear that the species is able to utilize carboxylates via a full TCA cycle and glyoxylate cycle. Multiple C4-dicarboxylate transporters, which are predicted to transport TCA cycle intermediates like malate, citrate, fumarate, and 2-oxoglutarate, are present as well. *Limnohabitans* sp. 1 may also be able to utilize fatty acids, as it encodes for the beta-oxidation pathway allowing for degredation of long-chain fatty acids. This is likely to occur, as *Chlamydomonas reinhardtii* secretes long-chain fatty acids (Jia, 2016). Another potential nutrient source for *Limnohabitans* sp. 1 is in sialic acid, for which multiple transporters are encoded (*siaM*, *siaQ*). However, the MAG does not encode for an enzyme to cleave sialic acids from host tissue, so it is unclear whether this is functional.

*Limnohabitans* sp. 1 is able to assimilate nitrogen directly from ammonia, an abundant nitrogen source in *Daphnia* (Hessen, 1990), via an NADP-specific glutamate dehydrogenase (*gdhA*), and encodes for an ammonia channel (*amtB*) to transport it directly into the bacterial cell.

*Biosynthesis pathways*

Synthesis of important molecules for cell structure and stability can be performed via gluconeogenesis, and *Limnohabitans* sp. 1 can begin this process via the conversion of malate to pyruvate via malic enzyme. The phospholipid biosynthesis pathway for production of phosphatidylethanolamine is present. Interestingly, the genome also encodes for the biosynthesis of dTDP-rhamnose, an important lipopolysaccharide production pathway (Gao, 2001). To synthesize nucleotides, *Limnohabitans* sp. 1 encodes for the PRPP (phosphoribosyl pyrophosphate) biosynthesis pathway from ribose-5-phosphate.

This MAG does not encode for biosynthesis pathways for all 21 amino acids. Full pathways are present for twelve amino acids: guanine from inosine; threonine, homoserine, and lysine from aspartate; cysteine from serine; valine from pyruvate; isoleucine from pyruvate or threonine; proline and arginine from glutamate via the orginine biosynthesis pathway; methionine from homocysteine; and phenylalanine and tyrosine from chorismate. Notably absent are pathways for histidine, taurine, tryptophan, valine, glutamine, glycine, and asparagine. A histidine degradation pathway and some evidence of a histidine transport system (*hisM*, *hisP*) are present, though a full transporter is not identifiable. The genome encodes for multiple amino acid transporters, including arginine and threonine exporters as well as cysteine and proline transporters. As there is no biosynthesis pathway present for glutamine, yet glutamine is shown to allow for *Limnohabitans* growth (Hornak, 2017), it is possible that that *Limnohabitans* sp. 1 relies on exogenous amino acids that it cannot synthesize. However, there is little evidence that amino acid degradation is occurring in this MAG; the only degradation pathway fully present is the histidine to glutamate pathway. The *Limnohabitans* sp. 1 arginine exporter (*argO*) and threonine exporter (*rhtA*) suggest that this species may be exporting essential amino acids for the host *Daphnia*.

This MAG encodes for some vitamin and cofactor biosynthesis pathways, but likely relies on other sources for many necessary molecules. Tetrahydrofolate, a folic acid derivative used in amino acid biosynthesis, can be synthesized. Glutathione, which is implicated in the reduction of ROS and detoxification of harmful molecules, can be biosynthesized. Pantothenate (vitamin B5) precursors can be transported into the bacteria via pantothenate precursor transporter PanS, then used to synthesize the essential cofactor coenzyme A. Transporters for vitamin B12 (cobalamin) and riboflavin (vitamin B2) are present.

*Host and environment interaction*

Adhesins, pili, and virulence factors are all implicated in bacterial colonization of a host through biofilm formation and host cell surface attachment. Secretion systems also are implicated in host cell invasion, as they can allow bacteria to transport proteins outside of the cell into the environment. It is likely that *Limnohabitans* uses adhesins to form biofilms, as it has been found to be a major constituent in aquatic biofilms (Ling, 2013), but because adhesin genes are extremely large it is unlikely that these would have been fully assembled in metagenome sequencing. Genes encoding for Type IV pili biogenesis (*pilQ*), genes involved in flagellar assembly (*flhA*, *flhB*, *flgB - L*), and virulence factors and regulators were found in the *Limnohabitans* sp. 1 MAG (*phoB-phoS*). Genes for chemotaxis were also present (*cheA*, *cheR*, *cheW*) and swarming motility (*swrC*, *rssA*).

The gene for microbial stealth protein CpsY, known to allow microbes to evade host immune defense systems, was identified in the genome (Sperisen, 2005). *Limnohabitans* sp. 1 encodes for a type 2 secretion system. RTX proteins and a type 1 secretion system membrane fusion protein (prsE) were also identified in this MAG, indicating that *Limnohabitans* sp. 1 may also have a type 1 secretion system. The genome also encodes for a Tat system, allowing for the transport of fully folded proteins across the cell membrane. Genes encoding a microcin C transporter were found, suggesting potential defense, detection, and regulation of microbial species and strains in the microbiota via microcins (Duquesne, 2007). An iron superoxide dismutase and catalase are present in this MAG as well, suggesting the ability to tolerate oxidative stress.

Quorum sensing is a way for bacterial species to communicate in response to their local environment. Genes involved in quorum sensing molecules can trigger functions involved in colonization and motility. *Limnohabitans* sp. 1 encodes for the QseB/QseC two-component regulatory system, which allows for the detection of signaling molecules. A putative gene (*ytnP*) involved in quorum quenching, which disrupts signaling between microbes and between microbes and the host, was identified in *Limnohabitans* sp. 1.

CRISPR-Cas systems allow bacteria to protect against foreign DNA. *Limnohabitans* sp. 1 contains multiple CRISPR-associated genes (*cas1*, *cas2*, *cas5d*, *cas9*). CRISPRCasFinder located 6 CRISPR elements with 49 spacer elements. The genome also contained genes for transposons (*tnpR*, *tnsB*, *tetD*) and prophage integrases (*intA*, *intS*), allowing for potential horizontal gene transfer.

*Summary*

Overall, *Limnohabitans* sp. 1 appears to be a mostly heterotrophic aerobe that primarily utilizes carboxylates and long-chain fatty acids to biosynthesize important molecules via the aerobic TCA cycle or the glyoxylate pathway. They may also be able to supplement their growth, and possibly benefit the host, by fixing carbon via bacteriochlorophyll *a* reaction centers. Furthermore, the presence of arginine and threonine exporters and the lack of host biosynthesis pathways for these two amino acids strongly indicates a benefit for the host by this bacterial species. Its wide range of potential substrates allows the species to be present at high abundance within the host *Daphnia* across multiple environments. The presence of microcin C transporters, type IV pili, and quorum-sensing systems suggest that *Limnohabitans* sp. 1 is able to detect and interact with other microbes in the *Daphnia* microbiota and with the host. The presence of bacteriochlorophyll suggests that this species is more likely to be highly abundant on the *Daphnia* carapace, rather than in the gut.

***Limnohabitans* sp. 2**

*Nutrient uptake and major energy pathways*

*Limnohabitans* sp. 2 contains genes encoding for a full cytochrome *c* oxidase, suggesting it is an aerobic species. Like sp. 1, *Limnohabitans* sp. 2 has the genetic potential to catabolize a wide range of nutrients. While the main glycolysis pathway is missing critical enzymes (hexokinase and phosphofructokinase), the full Entner-Doudoroff and the oxidative phase of the pentose phosphate pathway are present. It is likely that the full pentose phosphate pathway is present, as only one enzyme is missing (epimerase). This suggests that *Limnohabitans* sp. 2 is able to utilize glucose and other sugars as primary energy sources, further supported by the presence of a gene encoding a semiSWEET transporter. Transporters for alpha-glucosides are also present.

Alongside the Entner-Doudoroff pathway, *Limnohabitans* sp. 2 encodes for a complete TCA cycle. Like *Limnohabitans* sp. 1, this MAG contains genes encoding for multiple C4-dicarboxylate transporters predicted to transport the TCA cycle intermediates malate, fumarate, and 2-oxoglutarate, as well as succinate and aspartate with the C4-dicarboxylic acid transporter DauA. Uniquely encoded is a gentisate transporter (*genK*).

The full glyoxylate cycle is encoded in this MAG. In combination with the present of the two-component regulatory system for TCA transport (TctE-TctD), this suggests that *Limnohabitans* sp. 2 is able to process carbohydrates via multiple pathways for use in biosynthesis and cell maintenance. Other nutrient sources shared with *Limnohabitans* sp. 1 that may be utilized by this MAG include ammonia, sialic acid, and carbon fixed by use of bacteriochlorophyll a. Multiple sulfate transporters exist in the *Limnohabitans* sp. 2 MAG, but no complete sulfate reduction pathways are present. This suggests that *Limnohabitans* sp. 2 relies on other sources of sulfate, likely including cysteine.

*Biosynthesis pathways*

Like *Limnohabitans* sp. 1, sp. 2 does not encode for biosynthesis of all 21 amino acids. In fact, its amino acid biosynthesis capacity is limited to only leucine and arginine. Furthermore, the sp. 2 MAG does not contain basic purine or pyrimidine biosynthesis pathways in full. This lack of amino acid biosynthesis pathways may be partially caused by incomplete genome assembly, as *Limnohabitans* sp. 1 and sp. 2 are related and share 85% average nucleotide identiy. This is evidenced by the existence of multiple amino acid biosynthesis pathways with only one enzyme missing, including serine, threonine, betaine, cysteine, lysine, proline, histidine, tyrosine, and phenylalanine. More evidence of this is shown by the presence of a threonine/homeserine exporter (*rhtA*). It is also possible that some of the amino acids *Limnohabitans* sp. 2 is unable to synthesize are transported into the cell; importers and transporters for glutamate, cystine, proline, general branched-chain amino acids, and polar amino acids are present.

*Limnohabitans* sp. 2 can biosynthesize vitamin B6 using a pyridoxine-5-phosphate synthase (*pdxJ*) via pentose phosphate pathway intermediates, but relies on external sources for most of its other vitamins. A putative vitamin B1 (thiamine) transporter and transporters for tetrahydrofolate, glutathione, pantothenate, and cobalamin are present.

*Host and environment interaction*

Superoxide dismutase and catalase were found in the *Limnohabitans* sp. 2 MAG. Presence of the EnvZ-OmpR regulatory system indicates the ability to respond to changes in osmolality within *Daphnia* and in the environment. The CusS-CusR system is also present, which may allow *Limnohabitans* sp. 2 to tolerate high levels of copper. Genes involved in type 1, type 2, and Tat secretion were present (*prsE*, *gspF*, *epsE*, *outD*, *apxIB*, *tatA*, *tatB*, *tatC*). Two proteases associated with aminoglycoside resistance were also encoded (FtsH, HtpX). Overall, *Limnohabitans* sp. 2 is able to respond to multiple environmental stressors using different regulatory systems.

Microbial stealth protein CpsY, quorum-quenching lactonase YtnP, and the QseB/QseC quorum sensing regulatory system were also present. Genes for type IV pili and flagella were the same as found in *Limnohabitans* sp. 1; the only chemotaxis gene present was *cheR*. Swarming motility protein gene *swrC* and sensor protein gene *RssA* were present. Like *Limnohabitans* sp. 1, sp. 2 does not have any intact adhesin genes. However, a complete adhesin protein transport system is present. No CRISPR genes or transposon genes were found, making it unclear how the *Limnohabitans* sp. 2 MAG defends against foreign DNA. This MAG is the only one to encode for a heme export system (*ccmB*, *ccmC*) and heme binding proteins, which may indicate that this species is able to uptake and break down heme acquired from the host *Daphnia*.

*Summary*

Like sp.1, *Limnohabitans* sp. 2 is an aerobic species that can utilize carboxylates and carbohydrates. It is able to use glucose and other sugars via the Entner-Duodoroff pathway. *Limnohabitans* sp. 2 seems to be able to tolerate a wide range of environmental conditions due to its copper tolerance and osmotic regulatory system.

Its severely limited ability to biosynthesize amino acids, vitamins, and other important molecules suggests that *Limnohabitans* sp. 2 relies heavily on external, environmental sources for these molecules. Functional redundancy in the two *Limnohabitans* species, paired with the broader ability to function in a range of environmental conditions, suggests that sp. 2 is not as abundant as sp. 1 in the *Daphnia* microbiota under normal conditions but may be able to sustain necessary host-microbe functions in more stressful environmental conditions.

### Unknown Burkholderiaceae

#### *Nutrient uptake and major energy pathways*

The Burkholderia MAG contains genes encoding for NADH:quinone reductases and a cytochrome *bc1* complex. It also encodes for a full cytochrome *c* oxidase, suggesting it is a mostly aerobic species that can survive in low oxygen conditions if necessary. Important genes in glycolysis are not present. However, the Burkholderia genome encodes for the Leloir pathway, allowing for galactose to be metabolized to glucose. Methyl-galactoside may be imported into the cell via a transport system (*mglABC*), which can then be converted to galactose. The MAG also encodes for the conversion of oxaloacetate to fructose-6-phosphate as part of gluconeogenesis. The complete

TCA cycle and glyoxylate cycle are encoded as well, suggesting multiple methods of carbohydrate processing in the Burkholderiaceae genome. This is further evidenced by a maltose/trehalose transporter and a sorbitol/mannitol transporter, along with multiple C4-dicarboxylate transporters to import TCA cycle intermediates. A cation/acetate symporter allows acetate to be brought into a Burkholderiaceae cell; the two-step acetate kinase pathway allows for the production of acetyl-CoA from imported acetate.

The Burkholderiaceae species is able to utilize nitrate, importing it via the nitrite/nitrate transporter NarK. A respiratory nitrate reductase converts nitrate to nitrite for use in the cell, and NarK may be used to export overabundant nitrite. This dissimilatory nitrate reductase suggests the Burkholderiaceae uses this nitrate reduction pathway to accept electrons under low oxygen conditions. An Amt transporter exists to import extracellular ammonia. This extracellular ammonia can then be used to synthesize glutamate via glutamate dehydrogenase.

Though the Burkholderiaceae MAG contains different pathways for nitrogen use, some amino acid transporters still are identifiable. In particular, a proline/betaine transporter (*proP*) and a cystine transporter (*yijE*) are present, and components of the leucine-specific branched-chain amino acid transport system are present.

*Biosynthesis pathways*

The Burkholderiaceae MAG is able to synthesize purine and pyrimidine nucleotides from PRPP. Inosine monophosphate (IMP) is able to be synthesized from PRPP and glutamine, which is then used to synthesize both guanosine and adenosine. Uridine monophosphate (UMP) is able to be synthesized from the PRPP-biosynthesized orotate, which is then used to synthesize UTP and CTP. Complete pathways for fatty acid biosynthesis initiation and elongation were present.

This MAG does not encode for biosynthesis pathways for all 21 amino acids. Pathways for the biosynthesis of eleven amino acids are present: threonine, homeserine, cysteine, leucine, lysine, arginine, proline, tryptophan, phenylalanine, tyrosine, and glutathione. Pathways for the biosynthesis of serine, valine, isoleucine, and histidine were only missing one gene in the corresponding KEGG modules, suggesting they may be present. Pathways for the biosynthesis of taurine, glutamine, methionine, glycine, and asparagine were missing more than one gene. Like in *Limnohabitans* sp. 1, this MAG contained some evidence of a histidine transport (*hisP*, *his*Q), though the full transport system is not present. There is no indication of a glutamine transporter, but an ammonia transporter and a glutamine synthetase suggests that this species may be able to convert ammonia to glutamate. The genome encodes some amino acid transporters, including an arginine exporter (ArgO), a threonine-serine exchanger (SteT), and a proline-betaine transporter (ProP).

Few vitamin and cofactor biosynthesis pathways are present. Tetrahydrofolate, coenzyme A, and NAD biosynthesis pathways are complete. Vitamin B6 can be biosynthesized, and a cobalamin biosynthesis gene is present (*cbiB*).

*Host and environment interaction*

This MAG encodes for an osmotically-inducible protein (OsmY), which is part of an osmoprotectant ABC transporter complex. This protein likely helps to transport osmoprotectant molecules across the bacterial membrane, allowing the Burkholderiaceae species to survive under extreme osmotic stress. The BaeSR two-component system is present, allowing this MAG to respond to osmotic stress and some cell-damaging agents by upregulating efflux pump expression (Leblanc, 2011).

Superoxide dismutase, catalase, and two-component systems for the regulation of phosphate (CreCB) and nitrogen are present (GlnLG). The presence of nitrate respiration two-component system NarXL further suggests that the Burkholderiaceae MAG can utilize nitrogen when necessary.

Full pathways for a type IV secretion system, a general Sec secretion system, and the Tat secretion system are encoded in the MAG. Together, the Sec and Tat pathways may allow unfolded and folded proteins to be utilized by the type IV secretion system, which transports these proteins out of the cell into the environment or directly into other cells. Genes for movement through chemotaxis, swarming, or flagellar production were not identified in this MAG, nor were genes involved in biofilm production. However, multiple autotransporter adhesin genes were identified (*bmaC*, *ehaG*, *ata*), suggesting that cells are able to adhere to some host cell element. Like in both *Limnohabitans* MAGs, the QseB/QseC quorum sensing regulatory system was present. However, no quorum-suppressing or stealth protein genes were identified. No CRISPR genes or spacer genes were found, and only the prophage integrase *intS* gene was identified.

*Summary*

The Burkholderiaceae MAG is likely an aerobe that can survive in microaerophilic conditions by utilizing nitrogen. It has the potential to tolerate extreme osmotic stress. The *Daphnia* body and gut are mostly aerobic but the gut can have lower oxygen levels as food is digested (CITE THIS), suggesting that this species may live primarily in the lower gut of its host. It can uniquely utilize galactose as an energy source, and is able to synthesize most of its necessary amino acids. Presence of adhesin autotransporters and a lack of flagellar assembly genes and movement-related genes suggest that the species is able to directly adhere to the host *Daphnia* without moving. It is unclear if this Burkholderiaceae has any beneficial interactions with the host.

### *Pedobacter* sp.

#### *Nutrient uptake and major energy pathways*

This species contains genes encoding for cytochrome *c* oxidase and an NADH:quinone oxidoreductase, suggesting it is primarily an aerobe. It is also the only highly complete MAG to contain all genes necessary for glycolysis, suggesting it primarily uses glucose as an energy source. Glucose is able to enter the bacterial cell via a sodium/glucose cotransporter (*sglT*). *Pedobacter* can also biosynthesize glycogen from glucose-1-phosphate via the glgA, glgB, and glgC pathway. The *Pedobacter* MAG also encodes for a complete TCA cycle, and the same conversion of oxaloacetate to fructose-6-phosphate as the Burkholderiaceae MAG. Succinate and aspartate may be transported in as TCA cycle intermediates via DauA. An F-type ATPase is present as well.

Uniquely, *Pedobacter* is able to transport magnesium into the cell via the magnesium transporter MgtE. The *Pedobacter* MAG is also the only MAG to encode for a neuramindase, NanA. This may allow *Pedobacter* to cleave sialic acid from host cells; however, the *Pedobacter* MAG does not encode for a complete sialic acid transporter, so the presence of NanA may be to free sialic acid for other taxa that do encode for these transporters. In fact, both Limnohabitans MAGs reported here encode for sialic acid TRAP transporters but do not encode for a neuramindase, suggesting a potential interaction among these taxa.

*Biosynthesis pathways*

The non-oxidative phase of the pentose phosphate pathway can be used to biosynthesize ribose-5P, which can then be used to synthesize PRPP. From this, both purine and pyrimidine nucleotides can be produced. Like all of the other MAGs, *Pedobacter* does not encode for all amino acid biosynthesis pathways. In fact, it encodes for the fewest pathways: only isoleucine, leucine, tryptophan, and histidine can be produced. A nearly complete module for aspartate biosynthesis is also present. This is the only MAG to encode for histidine biosynthesis. Few biosynthesis pathways are fully encoded and there are no distinct amino acid transporters present, leaving it unclear how Pedobacter acquires necessary amino acids. One possible route of acquisition is through YhdG, a putative amino acid permease that allows amino acids to enter the cell.

Two unique isoprenoid biosynthesis pathways occur in the *Pedobacter* MAG, a C5 isoprenoid non-mevalonate pathway and a C10-C20 pathway. Isoprenoids are considered essential for pathogenic microbes (Odom, 2011), and these pathways may be mechanisms for *Daphnia* infection by *Pedobacter*.

*Pedobacter* can synthesize some cofactors and vitamins. The MAG encodes for the biosynthesis of NAD, a coenzyme for multiple redox reactions, from aspartate. Coenzyme A can be biosynthesized. Biotin, an essential cofactor for multiple processes in *Pedobacter*, can be biosynthesized from CoA and pimeloyl-ACP. Unlike the other MAGs, *Pedobacter* cannot biosynthesize tetrahydrofolate. It also cannot biosynthesize cobalamin or riboflavin.

*Host and environment interaction*

This MAG encodes for very few transporters overall. The only transporters present include a lipopolysaccharide transport system, a lipoprotein-releasing system, and a cell division transport system, indicating very little about environment interaction. Few permeases are present either, primarily consisting of amino acid permeases (*yhdG*) and macrolide export proteins (*macB*).

The *Pedobacter* MAG uniquely contains two chitinase A1 genes (*chiA*), suggesting that it may use chitin from algal food sources or from the *Daphnia* host. It also encodes a chitobiase (*chb*), allowing for degradation of chitin to N-acetylglucosamine. If *Pedobacter* is degrading chitin and cleaving sialic acid from the *Daphnia* host, this suggests that this species is not beneficial to the host. However, *Pedobacter* can biosynthesize biotin. Biotin is an essential vitamin for *Daphnia*, and growth media is typically supplemented with biotin to maximize host health. Harboring a species that produces biotin could be beneficial to the host in environments where there is little biotin available from food.

One persistence mechanism in *Pedobacter* may be through multidrug resistance. Both *Limnohabitans* MAGs and the *Polaromonas* MAG contain multiple genes encoding for antimicrobial agents, so other species may be able to persist if able to remove the antimicrobial agents using efflux pumps. *Pedobacter* encodes for the MdlAB/SmdAB transporter, a multidrug efflux pump that allows the cell to remove structurally different and complex antibiotics (Matsuo, 2008). There are also multiple macrolide export permease proteins encoded, suggesting that *Pedobacter* has multiple methods for antimicrobial agent removal from the cell.

The only secretion systems encoded in the *Pedobacter* genome are the Tat and Sec systems, suggesting they are able to secrete some folded and unfolded proteins into the local environment. Only the phosphate starvation response two-component system (PhoR-PhoB) is encoded here. CRISPR-associated genes Cas12a, Cas1, and Cas2 were identified. There were no transposon genes, or prophage integrase genes identified.

*Summary*

The *Pedobacter* species is likely an aerobe that persists on the carapace of *Daphnia magna*. Its ability to cleave sialic acid and dissolve chitin to digestible N-acetylglucosamine allow the species to utilize host cells for nutrients. The genome is not indicative of genes involved in movement, and few genes involved in nutrient transport appear to be present, suggesting that *Pedobacter* is able to persist on relatively few external nutrients. There are also almost no two-component regulatory systems present. The relatively high abundance of genes involved in antimicrobial efflux suggests that this *Pedobacter* MAG can persist in environments where other taxa, like the *Limnohabitans* species in *Daphnia*, are able to produce antimicrobial agents.

***Polaromonas* sp.**

*Nutrient uptake and major energy pathways*

The *Polaromonas* MAG contains genes encoding for a cytochrome *c* oxidase, a cytochrome *bc1* complex, and NADH:quinone oxidoreductases, suggesting it is similar to the unknown Burkholderiaceae in that it is an aerobe that can survive in low oxygen conditions. The complete TCA cycle is present, as is the glyoxylate cycle; however, multiple genes in glycolysis are not present.

*Polaromonas* is the only MAG in the *Daphnia* microbiota that may be sulfur oxidizing, as it contains a SOX complex and is able to oxidize thiosulfate to sulfate. Like in the Burkholderiaceae, sulfur oxidation may be a response to microaerophilic conditions. This is supported by the presence of a sulfate/thiosulfate transporter CysP and a sulfonate transport system SsuABC. Several amino acid transporters are also present, including a proline/betaine transporter (*proP*), a glutamate/aspartate importer (*gltK*), a cystine transporter (yijE), and a general branched-chain amino acid transporter.

*Biosynthesis pathways*

*Polaromonas* has the capability to biosynthesize both adenine and guanine, but does not have complete modules for pyrimidine ribonucleotide biosynthesis. The UMP to UDP/UTP biosynthesis pathway is only missing one block, suggesting that *Polaromonas* is able to synthesize pyrimidines. This is supported by the absence of nucleoside importers. Complete pathways for the biosynthesis of threonine, serine, cysteine, lysine, arginine, ornithine, proline, tryptophan, phenylalanine, and glutathione are present in this MAG. Betaine, isoleucine, histidine, and tyrosine biosynthesis pathways only have one block missing. With only the presence of a few amino acid importers, it appears that *Polaromonas* can biosynthesize the majority of its necessary amino acids.

Many important vitamins can be biosynthesized. Like all of the MAGs described above, *Polaromonas* can synthesize CoA from pantothenate. Like the *Pedobacter* species, it can synthesize biotin. There is no genomic evidence of biotin export. Tetrahydrofolate can be biosynthesized. Uniquely, *Polaromonas* can synthesize cobalamin (*cobA*, *cobQ*, *cbiB*, *cobP, cobC*) and riboflavin.

*Host and environment interaction*

As with the other MAGs, the *Poloromonas* genome encodes for superoxide dismutase and catalase. This MAG encodes for multiple transporters, many of which are shared with the *Limnohabitans* species. Shared transport systems include iron(III), sulfate, molybdate, tungstate, and nitrate/nitrate, among others included in Table S3. An Mla ABC transporter is encoded, allowing for removal of antibiotics and other toxic agents like gamma-hexachlorocyclohexane from the cell. This is further evidenced by the AcrAB-TolC/SmeDEF efflux pump, which confers resistance to many antibiotics and antimicrobial agents (Du, 2014). Another multidrug efflux pump, MdtAB, is also present.

The *Polaromonas* MAG encodes for multiple genes involved in movement and adhesion, including type IV biogenesis factors (*pilY1*, *pilQ*), twitching motility proteins (*pilT*), and fimbrial proteins (*pilE*). This MAG encodes for many genes involved in chemotaxis (*pctA*, *cheA*, *cheY*, *tar*, *cheB*, *mcp4*, *tsr*). The genome also encodes for a complete set of flagellar assembly machinery. The high number of genes involved in movement suggests that this is a highly mobile species.

Type I and II secretion genes are present, including those involved in the RaxAB-RaxC type I secretion system. The Tat system is also encoded. Multiple unique two-component systems are present, including the KdpD-KdpE system for regulating potassium transport, the AlgZ-AlgR system for alginate production, and the PilS-PilR type 4 fimbriae synthesis regulatory system. Other two-component regulatory systems present in the *Polaromonas* species and shared with other MAGs include PhoR-PhoB, QseC-QseB, CusS-CusR, GlnL-GlnG, and EnvZ-OmpR. Transposon genes are present (*tnsB*, *tetC*), as well as prophage integrase genes (*intA*, *intS*), but no CRISPR genes were annotated.

*Summary*

The *Polaromonas* MAG is likely an aerobe that can use thiosulfate to persist in microaerophilic environments. It has a relatively high number of genes involved in biofilm formation, movement, and flagellar assembly, suggesting it is mobile and could be able to travel within the *Daphnia* host. It is able to biosynthesize many of its necessary vitamins and most amino acids, and can uptake sialic acids via an ABC transporter. It can also export some amino acids, which may indicate a host interaction or an interaction with another microbial species within the host. It shares many features with the *Limnohabitans* species, suggesting it occupies a similar niche space within the host.

### Methods

### Supplementary results

*16S rRNA gene sequencing*

Sequencing of the V4 hypervariable region in samples of ultrapure water resulted in <50 read counts per sample, indicating little contamination of the DNA extraction kit or of reagents. In adult *D. magna*, six phyla were present: *Actinobacteria*, *Cyanobacteria*, *Bacteroidetes*, *Firmicutes*, *Proteobacteria*, and *Verrucomicrobia*. Cyanobacterial sequences identified in the *D. magna* samples were identical to those found in the *C. reinhardtii* samples, and are assumed to be chloroplast from the algae. In the remaining phyla within the adult *Daphnia*, the most prevalent bacterial families were the Burkholderiaceae and the Flavobacteriaceae. The COMBO media was similar to the *D. magna* microbiota in phyla present but families differed in relative abundance. Burkholderiaceae and Flavobacteriaceae relative abundance were greatly reduced in media, while other Bacteroidetes increased in abundance. Sphingobacteriaceae also had greater relative abundance in COMBO media than in *D. magna*.

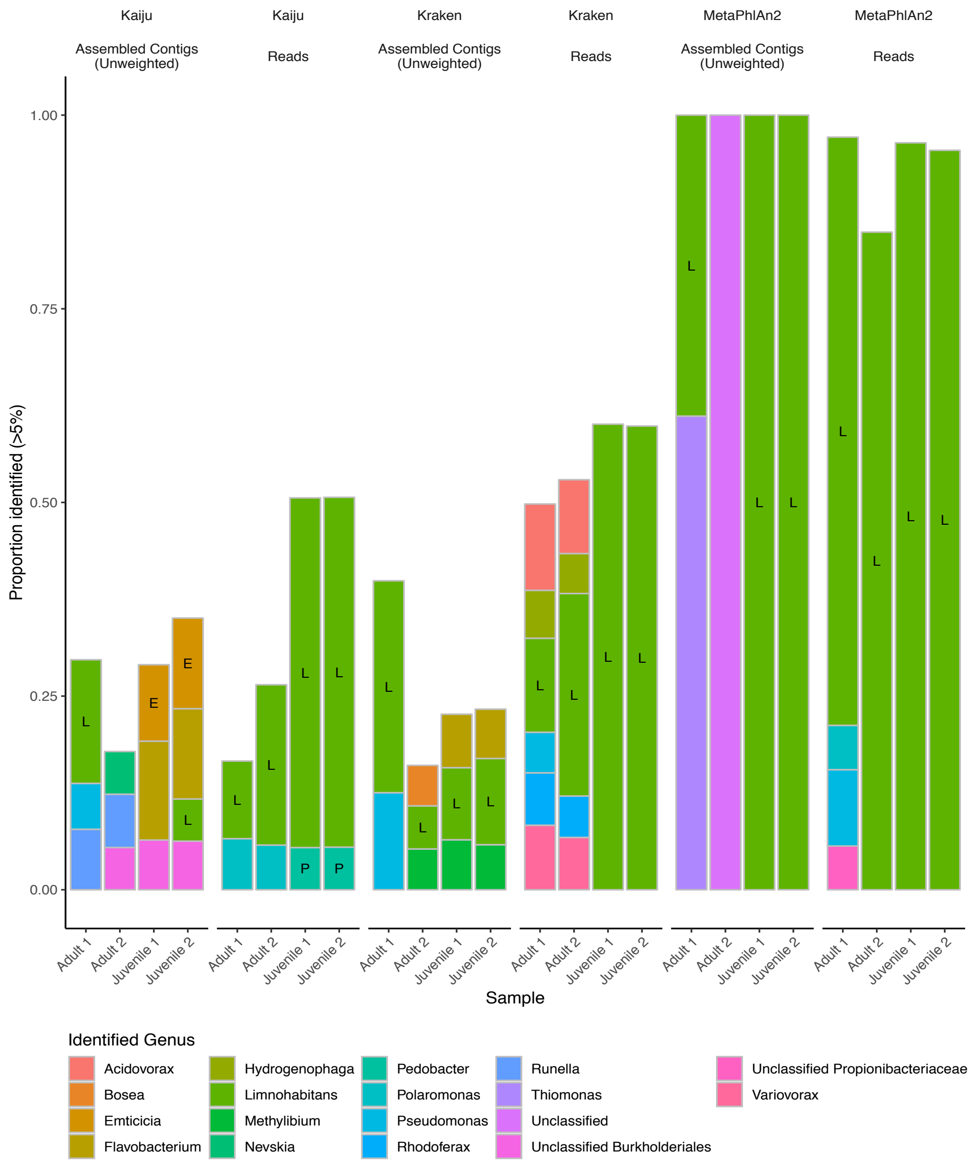
Kaiju
Assembled Contigs (Unweighted)
Kaiju
Reads
Kraken
Assembled Contigs (Unweighted)
Kraken
Reads
MetaPhlAn2
Assembled Contigs (Unweighted)
MetaPhlAn2
Reads
Proportion identified (>5%)
1.00
0.75
0.50
0.25
0.00
Adult 1
Adult 2
Juvenile 1
Juvenile 2
Sample
Identified Genus
Acidovorax
Bosea
Emticicia
Flavobacterium
Hydrogenophaga
Limnohabitans
Methylibium
Nevskia
Pedobacter
Polaromonas
Pseudomonas
Rhodoferax
Runella
Thiomonas
Unclassified
Unclassified Burkholderiales
Unclassified Propionibacteriaceae
Variovorax

**Supplementary Figure 1.** Taxonomic classification of reads and contigs from each sample using three taxonomic profiling tools: Kaiju, Kraken, and MetaPhlAn2. Bars represent genera with >5% relative abundance in read and contig data; letters within bars indicate genera identified through binning. All taxonomic profiling tools indicate the Daphnia magna microbiota is dominated by *Limnohabitans* species and that relative abundance differs by host life stage.

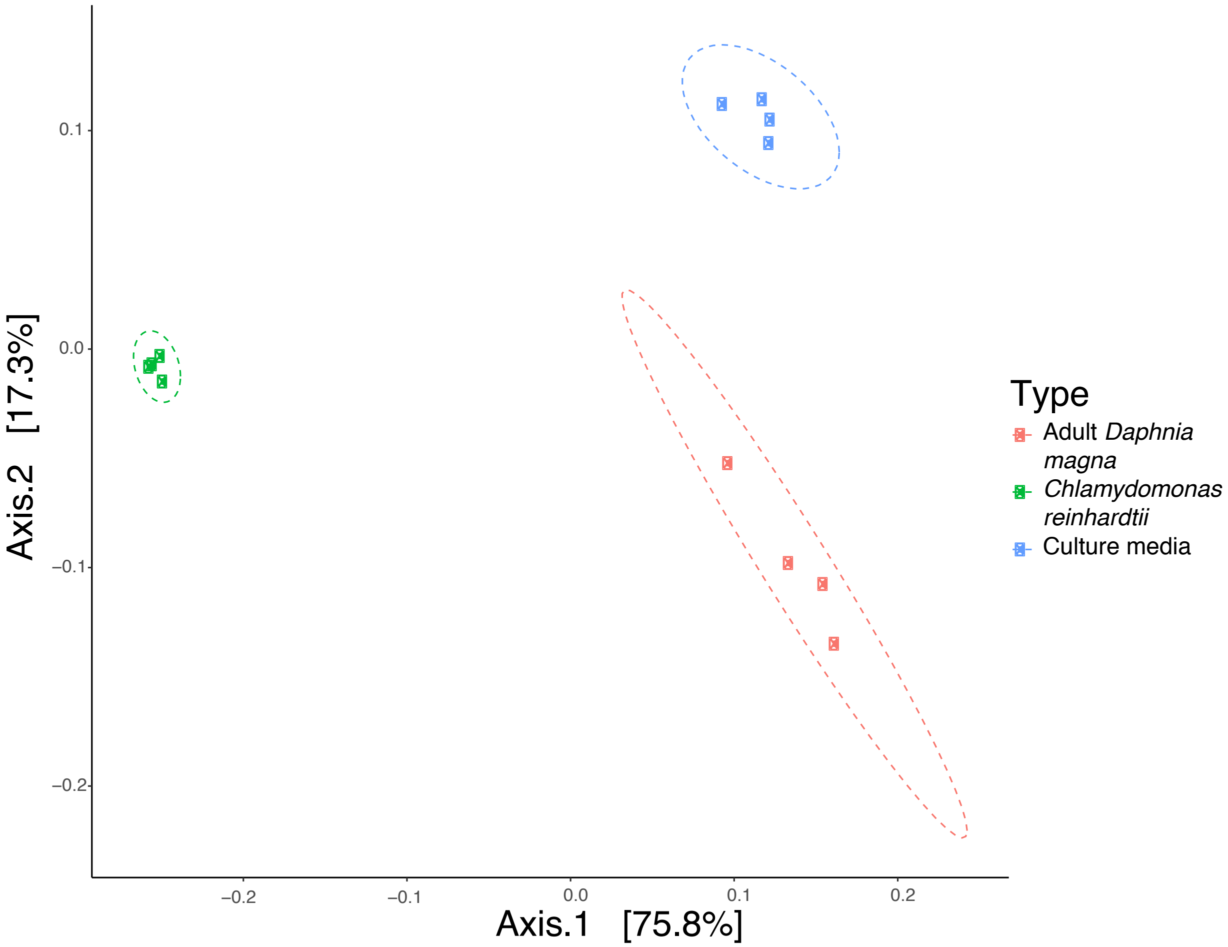

**Supplementary Figure 2.** PCoA plot of unweighted UniFrac distances between healthy adult *Daphnia magna* (red), the food source *Chlamydomonas reinhardtii* (green), and the COMBO medium used to culture *Daphnia magna* (blue). Each clusters independently.

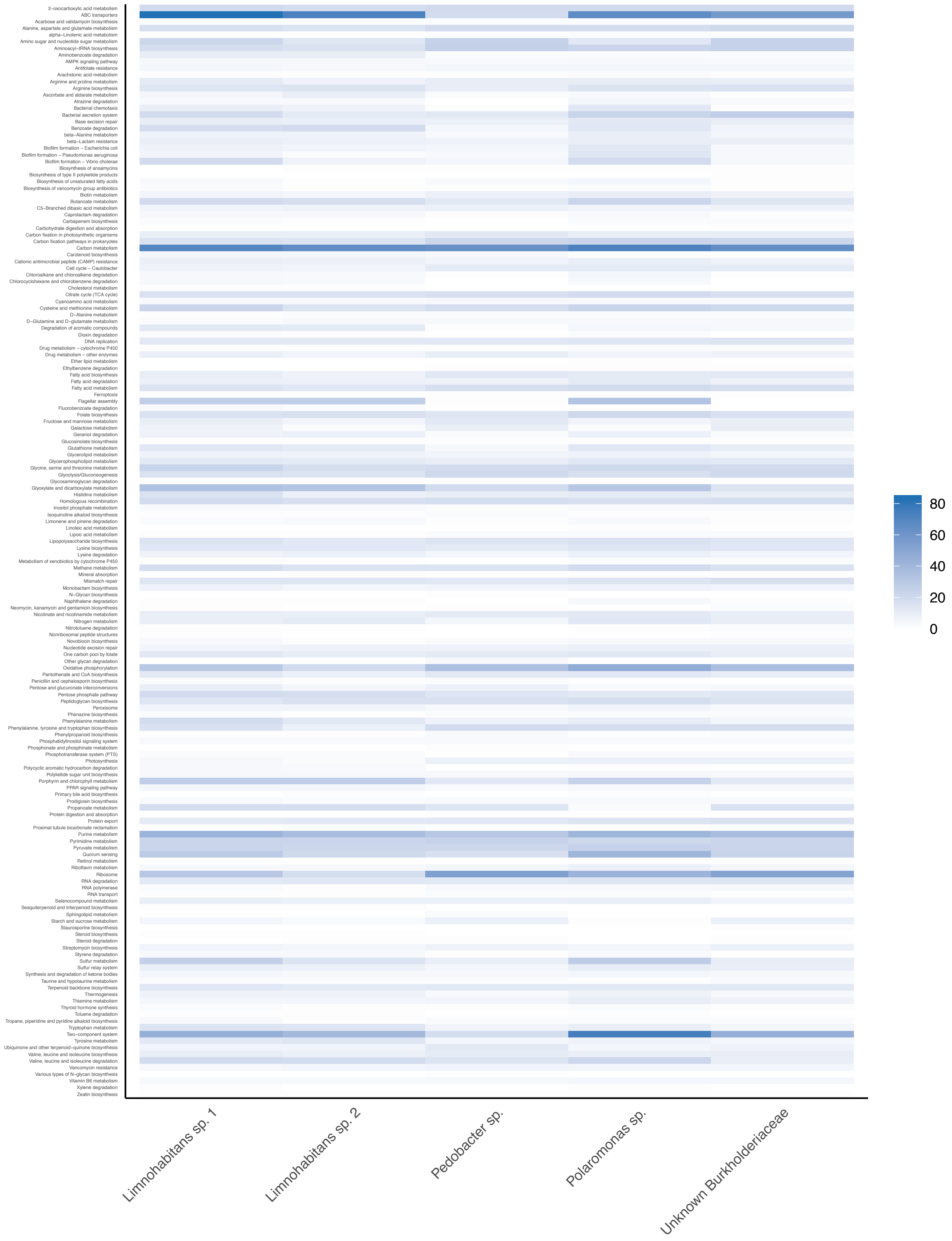
2–oxocarboxylic acid metabolism
ABC transporters
Acarbose and validamycin biosynthesis
Alanine, aspartate and glutamate metabolism
alpha–Linolenic acid metabolism
Amino sugar and nucleotide sugar metabolism
Aminoacyl–tRNA biosynthesis
Aminobenzoate degradation
AMPK signaling pathway
Antifolate resistance
Arachidonic acid metabolism
Arginine and proline metabolism
Arginine biosynthesis
Ascorbate and aldarate metabolism
Atrazine degradation
Bacterial chemotaxis
Bacterial secretion system
Base excision repair
Benzoate degradation
beta–Alanine metabolism
beta–Lactam resistance
Biofilm formation – Escherichia coli
Biofilm formation – Pseudomonas aeruginosa
Biofilm formation – Vibrio cholerae
Biosynthesis of ansamycins
Biosynthesis of type II polyketide products
Biosynthesis of unsaturated fatty acids
Biosynthesis of vancomycin group antibiotics
Biotin metabolism
Butanoate metabolism
C5–Branched dibasic acid metabolism
Caprolactam degradation
Carbapenem biosynthesis
Carbohydrate digestion and absorption
Carbon fixation in photosynthetic organisms
Carbon fixation pathways in prokaryotes
Carbon metabolism
Carotenoid biosynthesis
Cationic antimicrobial peptide (CAMP) resistance
Cell cycle – Caulobacter
Chloroalkane and chloroalkene degradation
Chlorocyclohexane and chlorobenzene degradation
Cholesterol metabolism
Citrate cycle (TCA cycle)
Cyanoamino acid metabolism
Cysteine and methionine metabolism
D–Alanine metabolism
D–Glutamine and D–glutamate metabolism
Degradation of aromatic compounds
Dioxin degradation
DNA replication
Drug metabolism – cytochrome P450
Drug metabolism – other enzymes
Ether lipid metabolism
Ethylbenzene degradation
Fatty acid biosynthesis
Fatty acid degradation
Fatty acid metabolism
Ferroptosis
Flagellar assembly
Fluorobenzoate degradation
Folate biosynthesis
Fructose and mannose metabolism
Galactose metabolism
Geraniol degradation
Glucosinolate biosynthesis
Glutathione metabolism
Glycerolipid metabolism
Glycerophospholipid metabolism
Glycine, serine and threonine metabolism
Glycolysis/Gluconeogenesis
Glycosaminoglycan degradation
Glyoxylate and dicarboxylate metabolism
Histidine metabolism
Homologous recombination
Inositol phosphate metabolism
Isoquinoline alkaloid biosynthesis
Limonene and pinene degradation
Linoleic acid metabolism
Lipoic acid metabolism
Lipopolysaccharide biosynthesis
Lysine biosynthesis
Lysine degradation
Metabolism of xenobiotics by cytochrome P450
Methane metabolism
Mineral absorption
Mismatch repair
Monobactam biosynthesis
N–Glycan biosynthesis
Naphthalene degradation
Neomycin, kanamycin and gentamicin biosynthesis
Nicotinate and nicotinamide metabolism
Nitrogen metabolism
Nitrotoluene degradation
Nonribosomal peptide structures
Novobiocin biosynthesis
Nucleotide excision repair
One carbon pool by folate
Other glycan degradation
Oxidative phosphorylation
Pantothenate and CoA biosynthesis
Penicillin and cephalosporin biosynthesis
Pentose and glucuronate interconversions
Pentose phosphate pathway
Peptidoglycan biosynthesis
Peroxisome
Phenazine biosynthesis
Phenylalanine metabolism
Phenylalanine, tyrosine and tryptophan biosynthesis
Phenylpropanoid biosynthesis
Phosphatidylinositol signaling system
Phosphonate and phosphinate metabolism
Phosphotransferase system (PTS)
Photosynthesis
Polycyclic aromatic hydrocarbon degradation
Polyketide sugar unit biosynthesis
Porphyrin and chlorophyll metabolism
PPAR signaling pathway
Primary bile acid biosynthesis
Prodigiosin biosynthesis
Propanoate metabolism
Protein digestion and absorption
Protein export
Proximal tubule bicarbonate reclamation
Purine metabolism
Pyrimidine metabolism
Pyruvate metabolism
Quorum sensing
Retinol metabolism
Riboflavin metabolism
Ribosome
RNA degradation
RNA polymerase
RNA transport
Selenocompound metabolism
Sesquiterpenoid and triterpenoid biosynthesis
Sphingolipid metabolism
Starch and sucrose metabolism
Staurosporine biosynthesis
Steroid biosynthesis
Steroid degradation
Streptomycin biosynthesis
Styrene degradation
Sulfur metabolism
Sulfur relay system
Synthesis and degradation of ketone bodies
Taurine and hypotaurine metabolism
Terpenoid backbone biosynthesis
Thermogenesis
Thiamine metabolism
Thyroid hormone synthesis
Toluene degradation
Tropane, piperidine and pyridine alkaloid biosynthesis
Tryptophan metabolism
Two–component system
Tyrosine metabolism
Ubiquinone and other terpenoid–quinone biosynthesis
Valine, leucine and isoleucine biosynthesis
Valine, leucine and isoleucine degradation
Vancomycin resistance
Various types of N–glycan biosynthesis
Vitamin B6 metabolism
Xylene degradation
Zeatin biosynthesis
Limnohabitans sp. 1
Limnohabitans sp. 2
Pedobacter sp.
Polaromonas sp.
Unknown Burkholderiaceae
80
60
40
20
0

**Supplementary Figure 3**. Heatmap of the number of genes identified in metabolic pathways using GHOSTKoala. The MAG with those genes is identified on the x-axis, and pathways are described on the y axis.

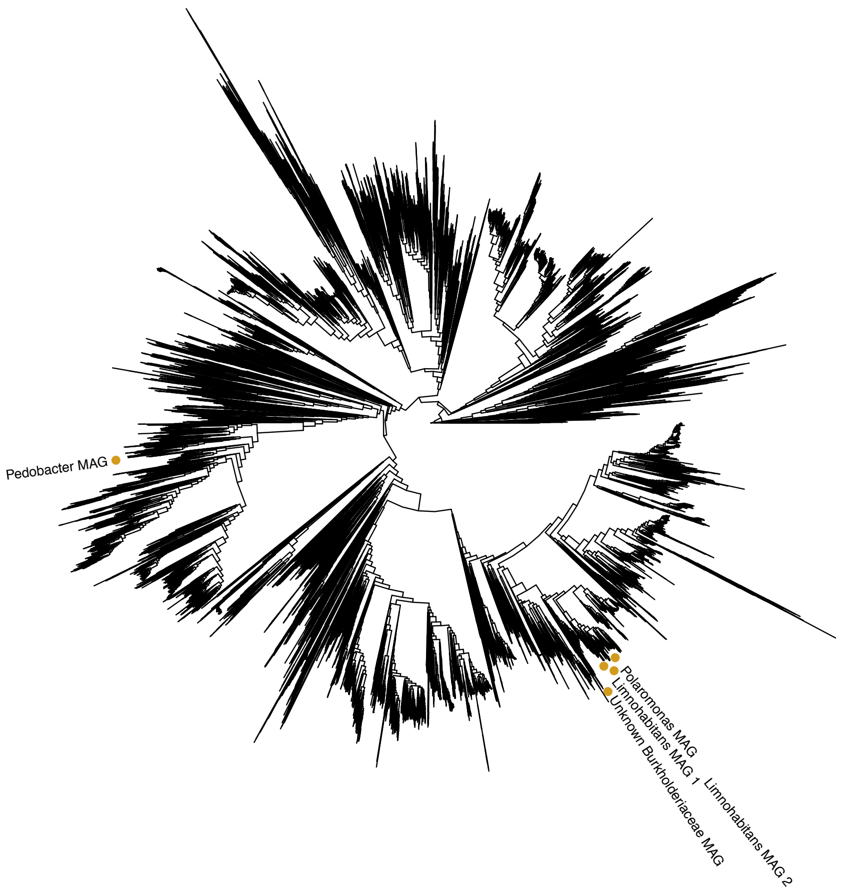
Pedobacter MAG
Polaromonas MAG
Limnohabitans MAG 1
Unknown Burkholderiaceae MAG
Limnohabitans MAG 2

**Supplementary Figure 4.** Placement of the five high- and medium-quality MAGs in the GTDB-Tk bacterial tree of life. Placement was performed using GTDB-Tk using pplacer and FastTree.

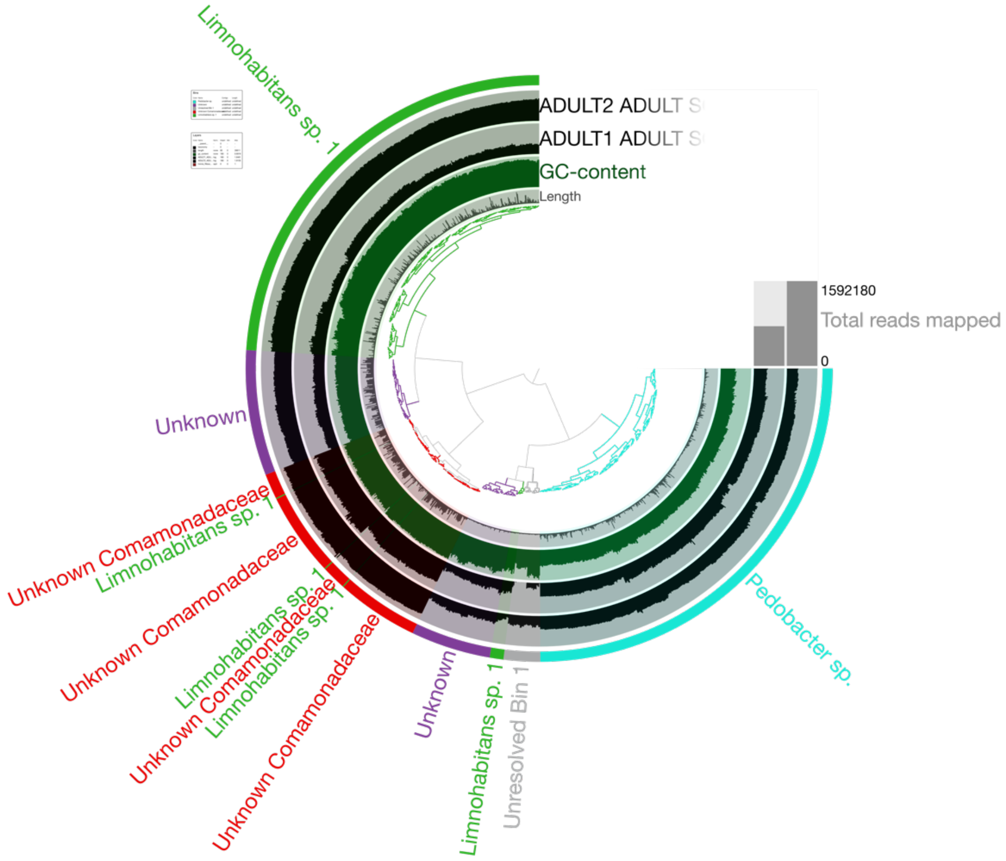

Limnohabitans sp. 1
ADULT2 ADULT S
ADULT1 ADULT S
GC-content
Length
1592180
Total reads mapped
0
Unknown
Unknown Comamonadaceae
Limnohabitans sp. 1
Unknown Comamonadaceae
Limnohabitans sp. 1
Unknown Comamonadaceae
Limnohabitans sp. 1
Unknown Comamonadaceae
Unknown
Limnohabitans sp. 1
Unresolved Bin 1
Pedobacter sp.

**Supplementary Figure 5.** Results of assembly and binning of the adult samples only represented in Anvi'o. Identification of bins was performed within Anvi'o using Centrifuge.

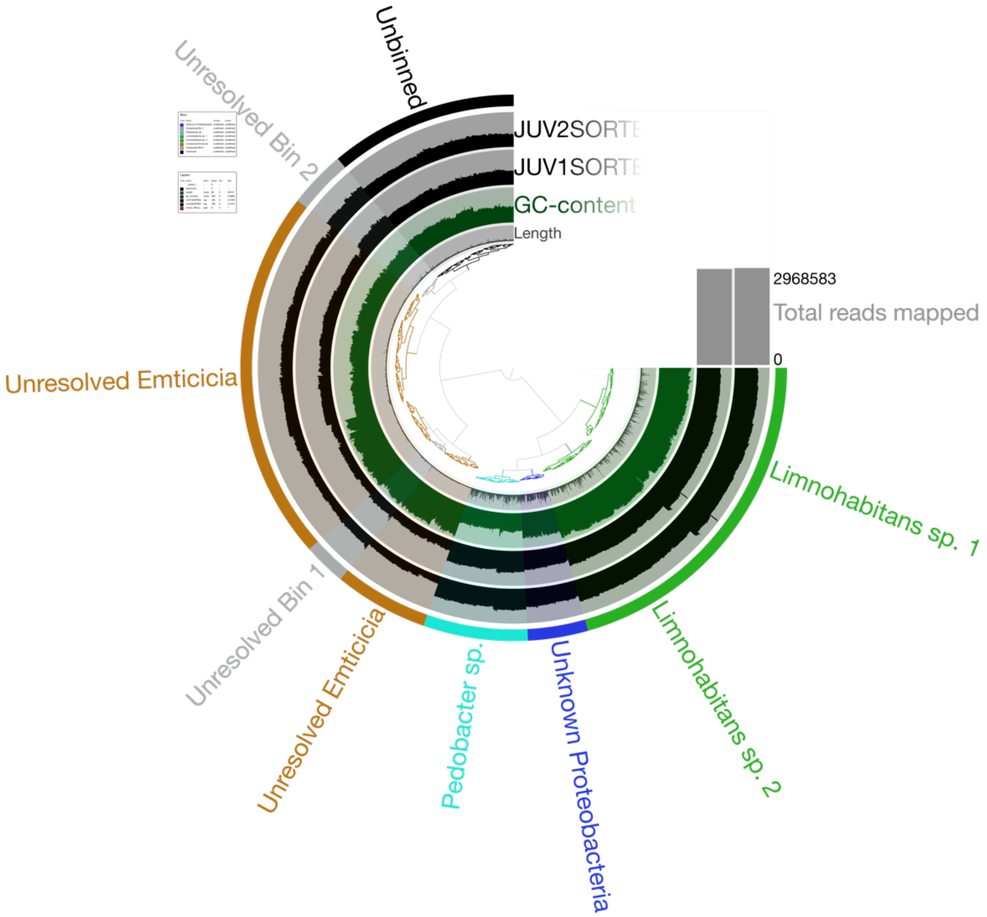
Unbinned
Unresolved Bin 2
JUV2SORTE
JUV1SORTE
GC-content
Length
2968583
Total reads mapped
0
Unresolved Emticicia
Limnohabitans sp. 1
Limnohabitans sp. 2
Unknown Proteobacteria
Pedobacter sp.
Unresolved Emticicia
Unresolved Bin 1

**Supplementary Figure 6.** Results of assembly and binning of the adult samples only represented in Anvi'o. Identification of bins was performed within Anvi'o using Centrifuge.

**Supplementary Table 1.** Summary of CONCOCT bins in the master co-assembly, the adult co-assembly, and the juvenile co-assembly. Identification of genomes was performed with Centrifuge within Anvi'o. Assessment of genome completeness is a measure of presence of single-copy gene sets.

**Supplementary Table 2**. Complete KEGG modules in each of the five high- and medium-quality MAGs. Dots represent that the pathway is present. Bold lettering in the pathway name indicate that it is a pathway unique to one MAG.

**Supplementary Table 3.** Summary of unique genes found in each MAG. This includes the putative gene name, Enzyme Commission number and COG identifier, as well as the product of that gene.

**Supplementary Table 4.** Summary of KEGG pathways in each MAG. Numbers correspond to the number of genes found in each MAG associated with that metabolic pathway in the KEGG database.
